# A type VII secretion system toxin functioning as a biofilm-specific intercellular signal in *Bacillus subtilis*

**DOI:** 10.1101/2023.08.02.551746

**Authors:** Kazuo Kobayashi

## Abstract

Biofilms are cooperative bacterial communities, in which diffusible signal-mediated quorum sensing coordinates the activities of member cells. However, limited diffusion in extracellular polymeric substance (EPS)-filled biofilms suggests existence of another intercellular signaling mechanism that is more adapted to the biofilm environment. Here, I demonstrate that the *yfjA* operon of *Bacillus subtilis* encodes one such signaling mechanism. The *yfjA* operon was induced by the two-component system DegSU, which controls biofilm development. Deleting the *yfjA* operon resulted in increased expression of DegSU-regulated genes under nutrient-rich conditions, hastening the onset of sporulation in biofilms. The *yfjA* operon encoded the type VII secretion system effector toxin YFJ in addition to its related proteins. YFJ toxin-mediated intercellular competition required EPS components which promote close cell-cell association, suggesting that the contact-dependent YFJ toxin system functions specifically in biofilms. The YFJ toxin also functioned as an intercellular signal that controlled the activity of DegSU. I propose that the YFJ toxin system is a biofilm-specific intercellular signaling mechanism that controls cell fate in biofilms in response to nutrient conditions and population size.

## Introduction

Intercellular signaling is vital for coordinating the activities and behaviors of multicellular systems, including bacterial biofilms. Biofilms are organized, multicellular communities of bacteria encased in a matrix of extracellular polymeric substances (EPSs) (Flemming & Wingender, 2010). In biofilms, bacterial cells cooperate with one another, for instance, by sharing extracellular products, while competing with deleterious and cheater strains (Burmølle *et al*, 2014; Hibbing *et al*, 2010). To adapt to the multicellular lifestyle in biofilms, many bacteria utilize a bacterial intercellular signaling mechanism known as quorum sensing (Davies *et al*, 1998; Hancock & Perego, 2004; Inhülsen *et al*, 2012; Schwartz *et al*, 2012; Vidal *et al*, 2015; Zhu & Mekalanos, 2003), which is mediated by the secretion and detection of small diffusible signal molecules, termed autoinducers (Miller & Bassler, 2001). In response to autoinducer accumulation, quorum sensing not only induces biofilm formation but also regulates cooperative and competitive bacterial interactions in biofilms (Abisado *et al*, 2018).

Biofilms consist of clusters of cells and EPSs, which generate three-dimensional structures. These structures can physically and biochemically hinder fluid flow and substance diffusion, indicating that quorum sensing may not function well in developed biofilms (Stewart, 2003). This reduced compound mobility has profound effects on cellular functions and even promotes biofilm-specific phenotypes. For example, EPSs physically and biochemically prevent some antibiotics and toxins from inhibiting cells within biofilms (Billings *et al*, 2013; Doroshenko *et al*, 2014, Tseng *et al*, 2013). In response, biofilm cells produce biofilm-specific antimicrobials that are not hindered by EPSs (Kobayashi, 2021; Kobayashi & Ikemoto, 2019; Oluyombo *et al*, 2019; Rendueles *et al*, 2014). Although quorum sensing depends on diffusible extracellular signals, the effect of the biofilm environment on quorum sensing remains unclear. This is partially because quorum sensing in biofilms has been mostly studied in the context of the thin biofilms produced by Gram-negative γ-proteobacteria (Davies *et al*, 1998; Inhülsen *et al*, 2012; Zhu & Mekalanos, 2003), which secrete highly diffusible molecules, N-acyl homoserine lactones as autoinducers (Miller & Bassler, 2001). However, unlike Gram-negative bacteria, Gram-positive bacteria secrete oligopeptides as autoinducers (Miller & Bassler, 2001), the diffusion of which is more likely influenced by physical and biochemical factors within biofilms. I therefore hypothesize that Gram-positive bacteria have another type of intercellular signaling mechanism that is more adapted to the biofilm environment.

The Gram-positive soil bacterium *Bacillus subtilis* forms thick biofilms, presenting as wrinkled colonies on solid media, and floating biofilms called pellicles, at the air-liquid interface in liquid media (Branda *et al*, 2001). These structures are composed of bundles of cell chains held together by EPSs, including exopolysaccharides and TasA amyloid fibers (Branda *et al*, 2006; Branda *et al*, 2001; Kobayashi, 2007a; Romero *et al*, 2010). Under laboratory conditions, biofilm formation by *B. subtilis* is usually followed by sporulation, and stress-resistant dormant spores predominantly are formed at the upper surface of the biofilm as nutrient levels decline (Branda *et al*, 2001, Vlamakis *et al*, 2008). *B. subtilis* secretes multiple oligopeptides as autoinducers, which directly and indirectly activate multiple two-component systems involved in biofilm formation (Kalamara *et al*, 2018). One of the two-component systems is DegS-DegU (hereafter referred to as DegSU), which induces approximately 50 genes and operons in biofilms (Kobayashi, 2007b), including gen.es required for biofilm formation (Gao *et al*, 2015; Hobley *et al*, 2013; Kiley & Stanley-Wall, 2010; Yu *et al*, 2016), those which encode extracellular degradative enzymes for nutrient acquisition (Kobayashi, 2007b; Marlow *et al*, 2014a), antimicrobial genes for — microbial competition (Kobayashi, 2015; Kobayashi, 2021; Kobayashi & Ikemoto, 2019; Mariappan *et al*, 2012; Tsuge *et al*, 1999; Xu *et al*, 2014), and many others of unknown function. Highly active DegSU also promotes sporulation in biofilms (Marlow *et al*, 2014b). Although its mechanism of activation remains unclear, DegSU is activated by the small protein DegQ (Kobayashi, 2007b). The moonlighting glycosyltransferase EpsE activates DegSU by inhibiting flagellar rotation (Blair *et al*, 2008; Cairns *et al*, 2013; Chan *et al*, 2014). Transcription of *degQ* and the *epsA* operon, including *epsE*, is induced by quorum sensing (Kalamara *et al*, 2018), indicating that DegSU activation depends on quorum sensing in biofilms.

Once DegSU is activated, phosphorylated DegU (DegU∼P) activates transcription of *degQ* and represses transcription of flagellar genes, indicating the likely presence of other layers of DegSU regulation (Amati *et al*, 2004; Msadek *et al*, 1991). After being activated in biofilms, DegSU may still require intercellular signaling to function. DegSU contributes to intercellular competition in biofilms by inducing various antimicrobials, including multiple LXG toxins, which are exported by the type VII secretion system (T7SS) (Kobayashi, 2015; Kobayashi, 2021; Kobayashi & Ikemoto, 2019; Mariappan *et al*, 2012; Tsuge *et al*, 1999; Xu *et al*, 2014). Producing multiple antimicrobials is likely advantageous in the biofilm environment, but may be wasteful in the absence of competitors. Indeed, a recent study reported that some bacteria upregulate antimicrobial production in response to competitors (Gonzalez *et al*, 2018; Lazzaro *et al*, 2017; Mavridou *et al*, 2018; Westhoff *et al*, 2021). I therefore speculate that *B. subtilis* possesses an additional intercellular signaling mechanism that modulates DegSU activity in biofilms depending on whether the cell is in the presence of competitor strains.

Here, I investigated the DegSU-regulated *yfjA* operon as a candidate encoding the intercellular signaling functions downstream of DegSU. My results suggest that the *yfjA* operon encodes a T7SS toxin and its related proteins, which control cell fate as a biofilm-specific intercellular signaling mechanism.

## Results

### Identification of the *yfjA* operon

I hypothesized that disrupting the hypothetical intercellular signaling mechanism would change DegSU activity, leading to altered colony biofilms morphology (Cairns *et al*, 2013; Chan *et al*, 2014; Kobayashi, 2015; Verhamme *et al*, 2007). Further, since this mechanism functions during the DegSU-active phase, it may be encoded by unknown DegSU-regulated genes. I previously constructed disruption mutants of almost all DegSU-regulated genes and operons and assayed their biofilm morphology in nutrient-rich biofilm medium 2×SGG (Kobayashi, 2007b). One of the disruption mutants harboring deletion of the entire *yfjA* operon (hereafter referred to as Δ*yfjA-*op) formed macroscopically wrinkled colony biofilms, which were morphologically different from those of the parental strain NCIB 3610 [hereafter referred to as the wild-type (WT) strain in this study] (Fig 1A). WT and Δ*yfjA-*op strains grew similarly in agitated culture, suggesting that the phenotype of the Δ*yfjA-*op mutant was not due to growth defects (Fig S1). The formation of macroscopic wrinkles on colony biofilms is often promoted by increased production of EPSs such as poly-γ-glutamate (Kobayashi, 2015; Morikawa *et al*, 2006; Yu *et al*, 2016), which is synthesized by products of the DegSU-regulated *pgsBCA-ywsC* operon. I therefore examined whether poly-γ-glutamate contributes to the Δ*yfjA-*op mutant phenotype. Disrupting *pgsC* reversed the colony morphology of the Δ*yfjA-*op strain, although the Δ*pgsC* mutation alone also altered colony morphology (Fig 1A). These results suggest that the Δ*yfjA-*op mutation increases DegSU-dependent poly-γ-glutamate production, thereby promoting macroscopic wrinkle formation on colony biofilms. Unlike in solid medium, the Δ*yfjA-*op mutation had no effect on the morphology of pellicle biofilms in liquid standing culture (Fig S2). I therefore investigated whether the *yfjA* operon encodes an unknown intercellular DegSU regulatory mechanism in colony biofilms.

**Figure 1.**
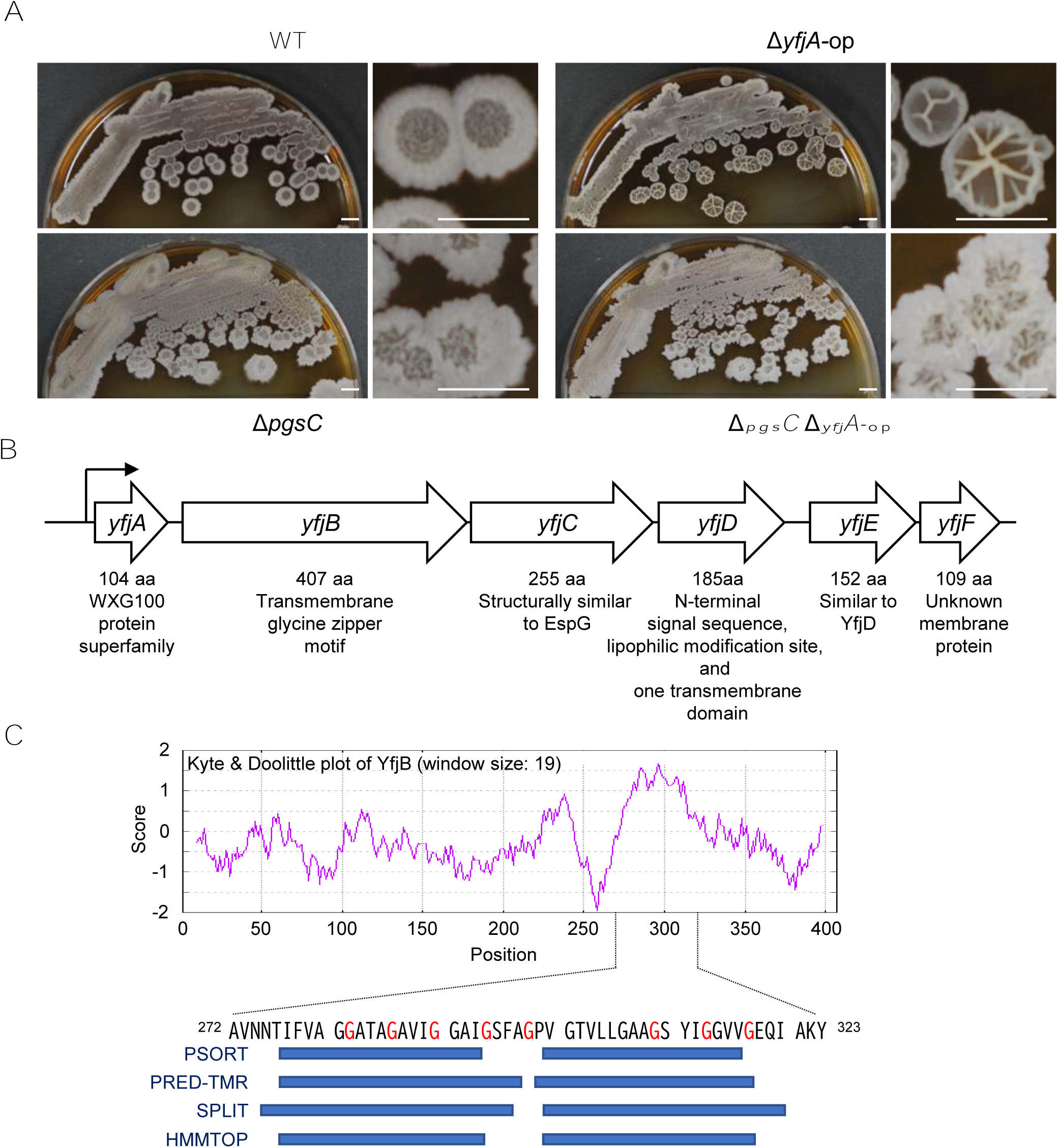
The *yfjA* operon encodes a candidate regulatory system for DegSU. A. Colony morphology of the Δ*yfjA-*op mutant. *B. subtilis* strains were grown in 2×SGG medium. Scale bar, 5 mm. B. Organization of the *yfjA* operon. Size and annotation of proteins encoded by individual genes are shown below the gene map. C. A Kyte & Doolittle hydropathy plot of YfjC. The window size is 19. Blue bars below the amino acid sequence indicate potential transmembrane regions predicted by PSORT (https://psort.hgc.jp/), PRED-TMR (http://athina.biol.uoa.gr/PRED-TMR/), SPLIT (http://split.pmfst.hr/split/4/), and HMMTOP (http://www.enzim.hu/hmmtop/index.php). Gly-xxx-Gly-xxx-Gly sequences are indicated by red G letters in the amino acid sequence.

The uncharacterized six gene operon *yfjABCDEF* is conserved among 13 fully sequenced *B. subtilis* strains, but is rarely found in other species (Fig S3). Each protein sequence was highly conserved among the *B. subtilis* strains (Fig S4). Sequence annotation provides some insight into the functions of *yfjA* operon products (Fig 1B), but none of these gene products appeared to regulate *degSU* as a DNA-binding transcriptional regulator. YfjA belongs to the WXG100 protein superfamily, members of which form heterodimers with T7SS substrates and facilitate substrate export as chaperones (Abdallah *et al*, 2007; Champion *et al*, 2006; Whitney *et al*, 2017). YfjB contains a hydrophobic region within its C-terminal half (Fig 1C). Although less hydrophobic than typical transmembrane domains with a Kyte & Doolittle score of ≥1.6 (Kyte & Doolittle, 1982), this region was bioinformatically predicted to form two transmembrane domains (Fig 1C). The predicted transmembrane domains contain GxxxGxxxG sequences, which match the transmembrane glycine zipper motif (Fig 1C). This motif is found in channel proteins and pore-forming bacteriocins and mediates their homo-oligomerization (Ali *et al*, 2023; Kim *et al*, 2004; Kim *et al*, 2005). YfjC had no significant sequence similarity to known proteins, but its predicted 3-D structure generated by AlphaFold (https://alphafold.ebi.ac.uk/entry/O31556) was similar to that of EspG chaperones of the mycobacterial T7SS (Tuukkanen *et al*, 2019) (Fig S5). The other proteins, YfjD, YfjE, and YfjF showed no significant similarity to known proteins. YfjD and YfjE themselves show degrees of sequence similarity, and encode a cleavable N-terminal signal sequence followed by a transmembrane domain and a large cytosolic sequence. Furthermore, YfjD, but not YfjE, contains a lipophilic modification site immediately downstream of the signal sequence. Unlike strain NCIB 3610 used in this study, many *B. subtilis* strains do not encode *yfjE* (Fig S3).

### Deletion of the *yfjA* operon increases transcription of DegSU-regulated genes

To confirm that DegSU induces the *yfjA* operon, I introduced a transcriptional fusion between the *yfjA* promoter and the green fluorescent protein (GFP) gene (hereafter referred to as P*_yfj_*_A_-*gfp*) to the chromosomal *amyE* locus and examined its expression in colony biofilms. Flow cytometry showed that the P*_yfj_*_A_-*gfp* strain produced stronger fluorescence than the background signal, and that the Δ*degU* mutation reduced the fluorescence intensity to background levels (Fig 2). These results support that DegSU induces the *yfjA* operon, either directly or indirectly.

**Figure 2.**
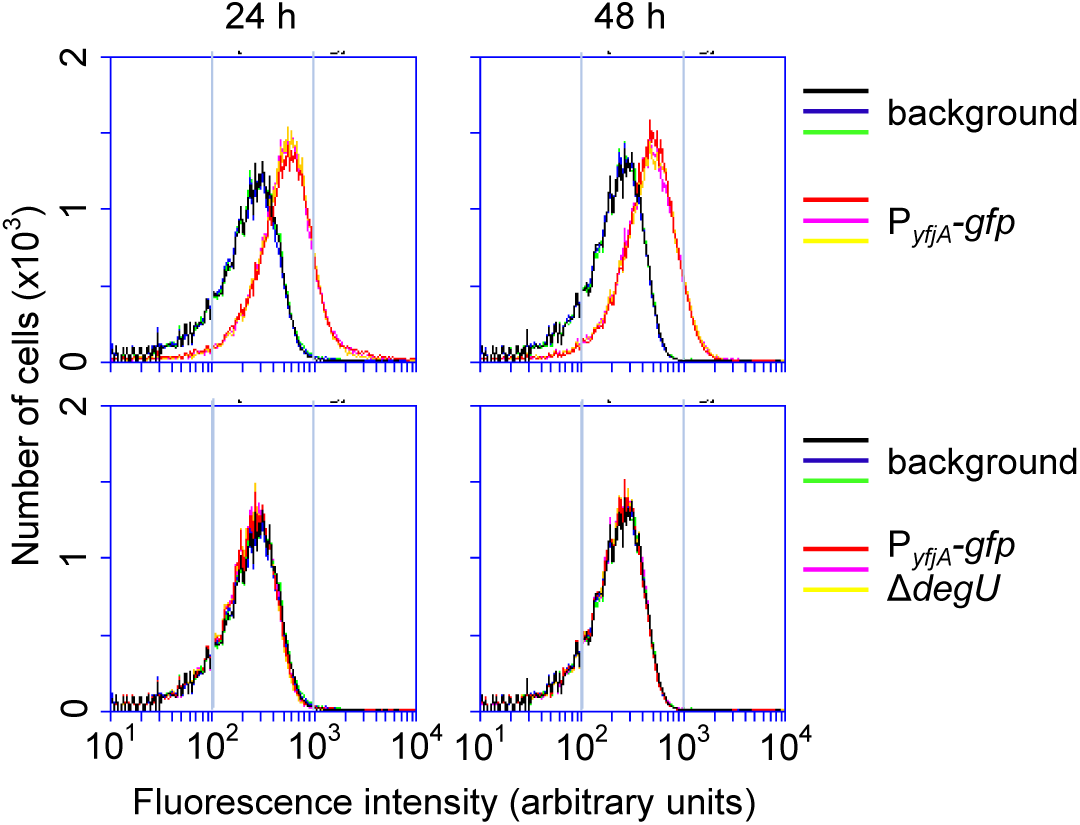
Expression of the *yfjA* operon. Strains harboring the P*_yfjA_*-*gfp* reporter were grown in 2×SGG medium. After 24 h or 48 h of cultivation, expression of *gfp* reporters in colonies was measured by flow cytometry. Strain 3610 (with no *gfp* reporter) was used as a background control. Three independent colonies were analyzed for each test.

To determine whether products of the *yfjA* operon regulate DegSU activity, I examined the effect of the Δ*yfjA-*op mutation on expression of the DegSU-regulated genes, *yfjA*, *bslA* (proteinaceous EPS) (Kobayashi & Iwano, 2012), and *aprE* (extracellular alkaline protease) (Mukai *et al*, 1990) using their respective promoter-*gfp* transcriptional fusions (Fig 3A). Flow cytometry analysis of the *gfp* reporter strains showed that the Δ*yfjA-*op mutant exhibited stronger fluorescence from P*_yfjA_*-*gfp* and P*_bslA_*-*gfp* than the WT strain. Median fluorescence values from reporters were 1.4-fold higher in the Δ*yfjA-*op mutant than in the WT strain (n=3). The Δ*yfjA-*op mutant had twice as many cells exhibiting strong fluorescence from P*_aprE_*-*gfp* as the WT strain. To quantify these changes, I more precisely compared GFP levels between P*_bslA_*-*gfp* and P*_bslA_*-*gfp* Δ*yfjA-*op strains. Since the fluorescence of GFP is unaffected by ≤0.5% SDS (Saeed & Ashraf, 2009), crude protein extracts of 24 h-old colony biofilms were separated by SDS-PAGE before fluorescent GFP bands in the gel were detected using a fluorescence imaging system. Quantitative analysis revealed that the Δ*yfjA-*op mutation increased GFP fluorescence from P*_bslA_*-*gfp* by 3.5-fold (Fig S6).

**Figure 3.**
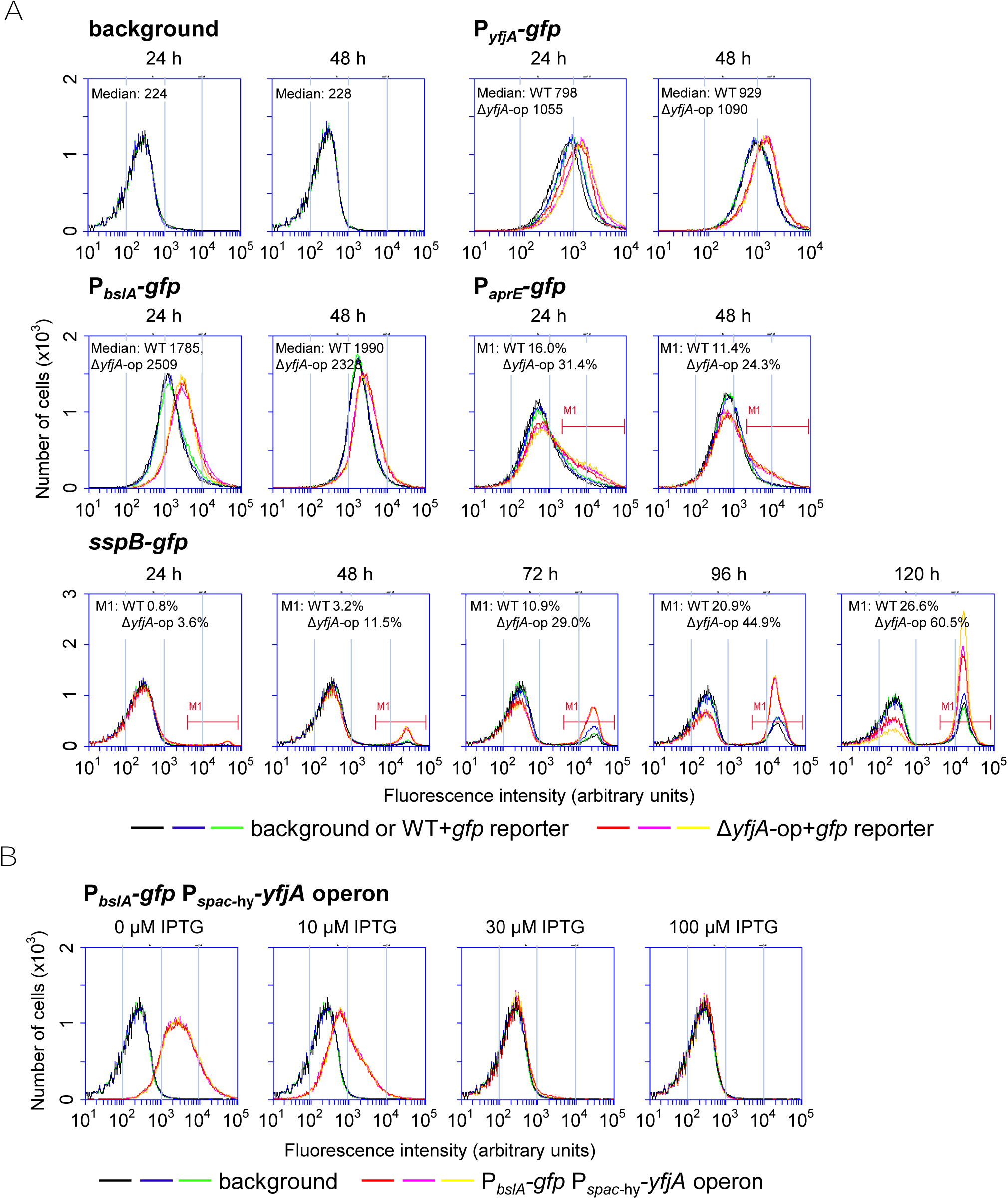
The Δ*yfjA-*op mutation increases expression of DegSU-regulated genes. A. The effect of Δ*yfjA-*op mutation on expression of DegSU-regulated genes. Strains harboring the indicated *gfp* reporters were grown in 2×SGG medium, and expression of *gfp* reporters was measured by flow cytometry. Three independent colonies were analyzed for each test. Median values of fluorescence peaks or percentages of population exhibiting strong fluorescence (M1) are indicated within the plots. B. Overexpression of the *yfjA* operon prevents *bslA* expression. The P*_spac_*_-hy_-*yfjA* operon strain harboring P*_bslA_*-*gfp* was grown in 2×SGG medium supplemented with indicated concentrations of IPTG, and expression of P*_bslA_*-*gfp* was measured at 24 h.

To verify that *yfjA* operon products negatively regulate DegSU, I overexpressed *yfjA* under the control of the isopropyl β-D-thiogalactopyranoside (IPTG)-inducible *spac*-hy promoter (a stronger variant of the IPTG-inducible *spac* promoter) (Quisel *et al*, 2001; Yansura & Henner, 1984) before assaying *bslA* expression. Induction of the *yfjA* operon from the *spac*-hy promoter with ≥30 µM IPTG completely inhibited *bslA* expression (Fig 3B). These results indicate that *yfjA* operon products negatively regulate DegSU activity.

### The Δ*yfjA-*op mutation promotes sporulation

Marlow *et al* (2014b) suggested that highly active DegSU promotes sporulation by increasing Spo0A∼P levels. I therefore examined the effect of the Δ*yfjA-*op mutation on sporulation using the *sspB*-*gfp* fusion gene (Nishikawa & Kobayashi, 2021). Since the small acid-soluble spore protein SspB is induced in the late stage of sporulation and coats a spore chromosome (Errington, 2003), the percentage of cells expressing the SspB-GFP fusion protein are a proxy for sporulation frequency. Flow cytometry showed that detectable GFP fluorescence from *sspB-gfp* began to appear at 72 h in the WT strain and at 48 h in the Δ*yfjA-*op mutant (Fig 3A). By 120 h, the population exhibiting GFP fluorescence increased to 26.6 % in the WT strain and 60.5% in the Δ*yfjA-*op mutant (n=3), indicating that the Δ*yfjA-*op mutation hastens and promotes sporulation.

I next determined whether the Δ*yfjA-*op mutation promotes sporulation through DegSU. Since DegSU activity can be artificially controlled by altering DegU levels (Marlow *et al*, 2014b; Verhamme *et al*, 2007), I placed *degU* under control of the IPTG-inducible *spac* promoter (strain hereafter referred to as P*_spac_-degU*) (Yansura & Henner, 1984) and compared the effect of its induction on *sspB-gfp* expression between WT and Δ*yfjA-*op backgrounds by fluorescence microscopy. Δ*yfjA-*op colonies exhibited earlier and more intense GFP fluorescence than WT colonies, similar to that observed by flow cytometry (Fig 4). In colonies of P*_spac_-degU* and P*_spac_-degU* Δ*yfjA-*op strains, GFP fluorescence appeared earlier and was more intense across 96 h of cultivation, as the IPTG concentration increased (Fig 4). Irrespective of IPTG concentration, GFP fluorescence consistently appeared earlier and with greater intensity in the P*_spac_-degU* Δ*yfjA-*op strain than in the P*_spac_-degU* strain. The effect of the Δ*yfjA-*op mutation on *sspB-gfp* expression was more pronounced when *degU* was strongly induced by high concentrations of IPTG, consistent with the stimulatory effect of the Δ*yfjA-*op mutation on DegSU. These data show that the Δ*yfjA-*op mutation moderately increases DegSU activity, which is sufficient to induce earlier and more potent sporulation in colony biofilms.

**Figure 4.**
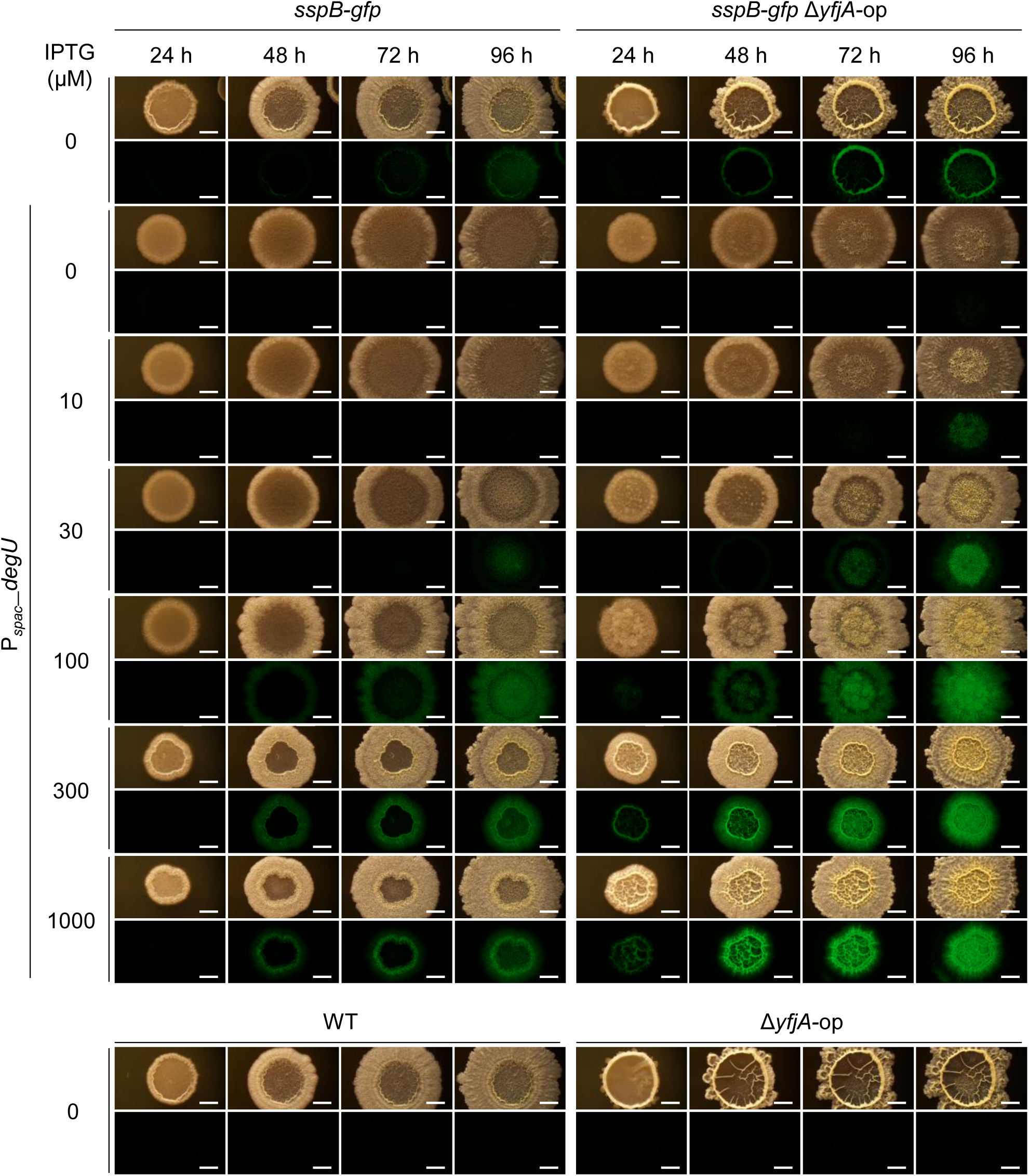
The Δ*yfjA-*op mutation promotes *sspB-gfp* expression through DegSU. *B. subtilis* strains were grown in 2×SGG solid medium supplemented with indicated concentrations of IPTG. Expression of *sspB-gfp* in colonies was analyzed by fluorescence microscopy every 24 h. Brightfield images and GFP images are shown. Images of WT and Δ*yfjA*-op strains without *sspB-gfp* are shown as negative controls. Scale bar, 2 mm.

### The phenotype of the Δ*yfjA-*op mutant depends on medium conditions

Unlike in the nutrient-rich biofilm medium 2×SGG, the Δ*yfjA-*op mutation had little or no effect on colony morphology and expression of *yfjA*, *bslA*, *aprE*, and *sspB* in the minimal biofilm medium MSgg (Branda *et al*, 2001) (Fig 5A, B). Indeed, flow cytometry analysis revealed that *yfjA*, *bslA*, and *aprE* were more strongly expressed in 2×SGG medium than in MSgg medium (Figs 3A and 5B). Similar expression patterns were previously observed in six other DegSU-regulated genes (Kobayashi, 2021). Moreover, Yu *et al*. (2016) previously reported that the DegSU-regulated *pgsB* operon was expressed in nutrient-rich complex medium but not in minimal medium. These observations indicate that DegSU is more strongly activated in nutrient-rich media than in minimal media.

**Figure 5.**
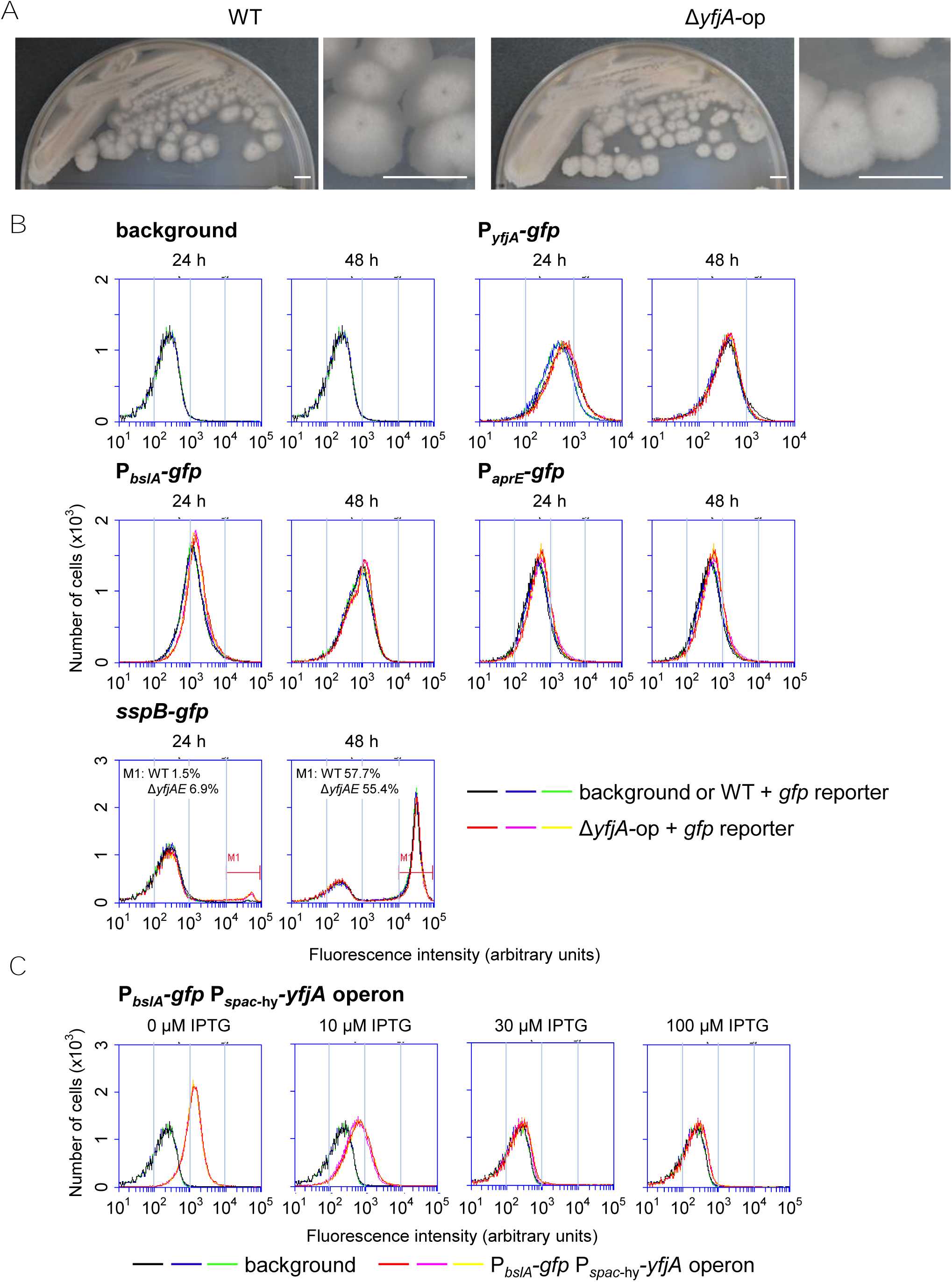
The Δ*yfjA-*op mutation has little or no effect on expression of DegSU-regulated genes in MSgg medium. A. Colony morphology of the Δ*yfjA-*op mutant. WT and Δ*yfjA-*op strains were grown for 72 h in MSgg medium. Scale bar, 5 mm. B. The effect of the Δ*yfjA-*op mutation on expression of DegSU-regulated genes in MSgg. After 24 h or 48 h of incubation, expression of *gfp* reporters in colonies were measured by flow cytometry. Strain 3610 (with no *gfp* reporter) was used as a background control. C. Overexpression of the *yfjA* operon prevents *bslA* expression in MSgg medium. The P*_spac_*_-hy_-*yfjA* operon strain harboring P*_bslA_*-*gfp* was grown in MSgg medium supplemented with indicated concentrations of IPTG, and expression of P*_bslA_*-*gfp* was measured at 24 h.

*yfjA* operon products retained DegSU inhibitory activity in MSgg medium as overexpression of the *yfjA* operon also inhibited *bslA* expression in MSgg (Fig 5C). I therefore hypothesized that *yfjA* operon products only exert an inhibitory effect on DegSU when highly active DegSU induces the *yfjA* operon. To test this hypothesis, I induced *degU* from the *spac* promoter with various concentrations of IPTG and compared *bslA* expression levels between the WT and Δ*yfjA-*op backgrounds. In P*_spac_*-*degU* and P*_spac_*-*degU* Δ*yfjA-*op strains, *bslA* expression gradually increased with increasing IPTG concentrations and almost reached WT levels at 100 µM IPTG (Fig 6). In the presence of ≤30 µM IPTG, *bslA* was similarly expressed in both strains, whereas in the presence of ≥100 µM IPTG, *bslA* was expressed more strongly in the P*_spac_*-*degU* Δ*yfjA-*op strain than in the P*_spac_*-*degU* strain. These results support the hypothesis.

**Figure 6.**
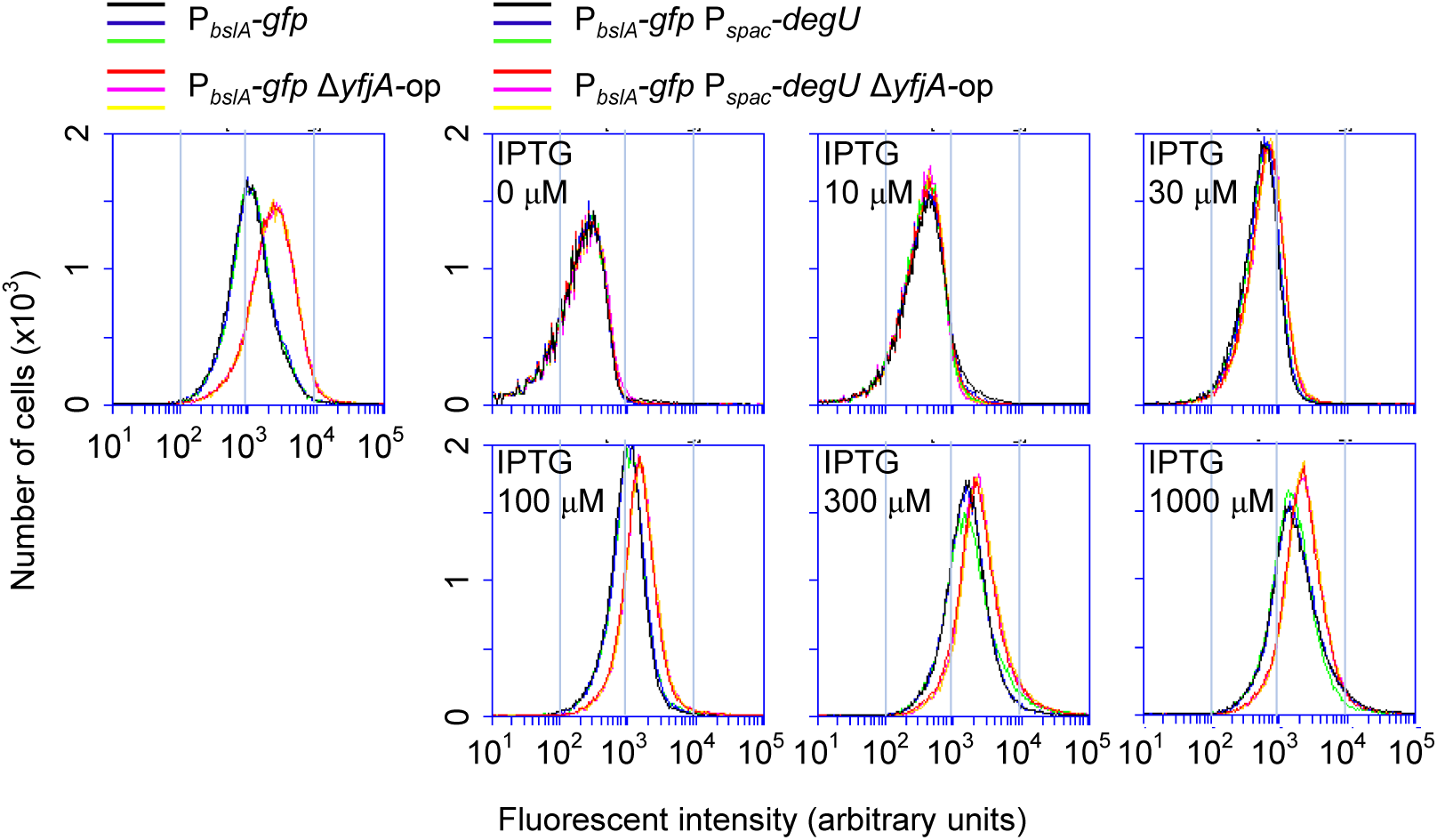
Functional expression of the *yfjA* operon requires high levels of DegSU. P*_spac_-degU* and P*_spac_-degU* Δ*yfjA-*op strains harboring P*_bslA_-gfp* were grown in 2×SGG medium supplemented with indicated concentrations of IPTG, and *bslA* expression was measured at 24 h. *bslA* expression in WT and Δ*yfjA-*op are shown as references.

The difference in *bslA* expression between these strains at ≥100 µM IPTG was smaller than that between WT and Δ*yfjA-*op strains. This was because at 300 µM and 1000 µM IPTG, the P*_spac_*-*degU* strain expressed more *bslA* than the WT strain, whereas the P*_spac_*-*degU* Δ*yfjA-*op strain expressed *bslA* at levels comparable to the Δ*yfjA-*op strain. Thus, induction of *degU* from the *spac* promoter appeared to dampen the effect of *yfjA* operon products. Since DegU∼P induces *degU* transcription (Kobayashi, 2007b; Ogura & Tsukahara 2010), the inhibitory effect of *yfjA* operon products on DegSU activity is expected to be enhanced by decelerating the positive autoregulation of *degU* in the WT strain. This suggests that the absence of the autoregulation in the P*_spac_*-*degU* strain can weaken the effect of the Δ*yfjA-*op mutation on DegSU activity, leading to a reduced difference in *bslA* expression between P*_spac_*-*degU* and P*_spac_*-*degU* Δ*yfjA-*op strains.

These data indicate that *yfjA* operon products exert an inhibitory effect on DegSU in a medium-dependent manner. Under nutrient-rich conditions, highly active DegSU strongly induces the *yfjA* operon, products of which prevent further activation of DegSU and delay the onset of sporulation.

### Identifying the function of individual *yfjA* operon genes

To investigate the function of individual genes of the *yfjA* operon, I constructed marker-less deletion mutants of *yfjA*, *yfjB*, and *yfjC*, and pMutin-vector-insertion mutants of *yfjD*, *yfjE*, and *yfjF.* The pMutin vector contains the IPTG-inducible *spac* promoter, which can drive transcription of genes located downstream of the insertion site (Vagner *et al*, 1998). I subsequently examined effects of these mutations on colony morphology and *bslA* expression (Fig 7A, B). Only Δ*yfjB* and Δ*yfjC* mutants formed macroscopic wrinkles on colonies and exhibited increased expression of *bslA* comparable to the Δ*yfjA*-op mutant.

**Figure 7.**
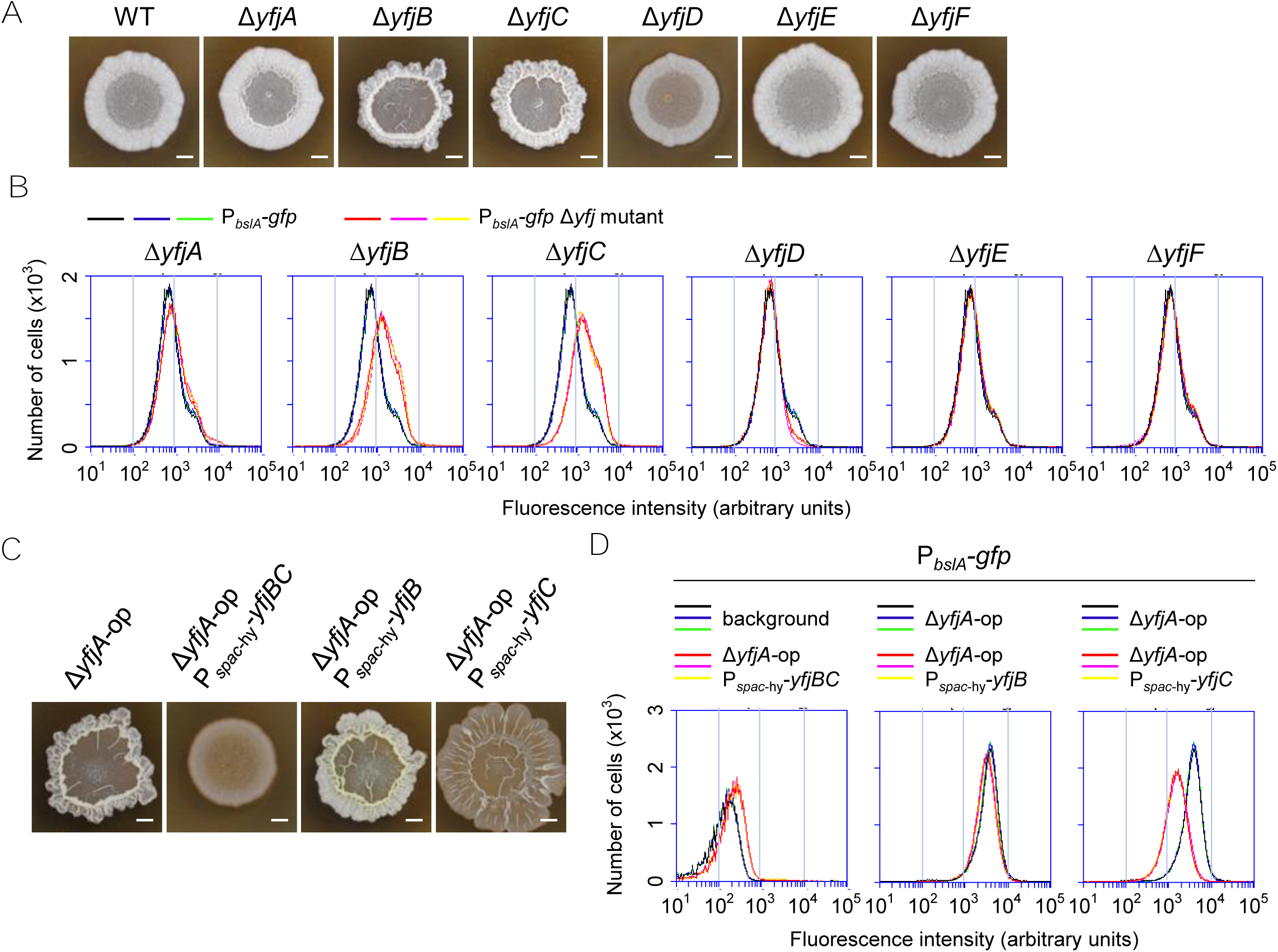
Functional analysis of individual *yfjA* operon genes. A. The colony morphology of *yfjA* operon gene mutants. Marker-less deletion mutants of *yfjA*, *yfjB* and *yfjC* were grown for 48 h in 2×SGG medium, and pMutin-insertion mutants of *yfjD*, *yfjE*, and *yfjF* were grown in 2×SGG supplemented with 100 μM IPTG. Scale bar; 2 mm. B. Expression of P*_bslA_-gfp* at 24 h in *yfjA* operon gene mutants. *bslA* expression in WT is shown as a reference. C. Overexpression of *yfjBC* prevents colony biofilm development. Strains were grown for 48 h in 2×SGG supplemented with 100 µM IPTG. Scale bar; 2 mm. D. Overexpression of *yfjBC* prevents *bslA* expression. Strains harboring P*_bslA_-gfp* were grown in 2×SGG supplemented with 100 µM IPTG and *bslA* expression was measured at 24 h. Background or *bslA* expression in of Δ*yfjA-*op are shown as references.

To test whether YfjB and YfjC are responsible for DegSU inhibition, I ectopically expressed *yfjB* and/or *yfjC* from the *spac*-hy promoter in the Δ*yfjA-*op mutant and examined their effects on colony morphology and *bslA* expression using the P*_bslA_-gfp* transcriptional fusion inserted into the chromosomal *htpG* locus. Overexpression of *yfjBC* completely inhibited colony wrinkle formation and *bslA* expression (Fig 7C, D). By contrast, overexpression of *yfjB* had no effect on colony morphology and *bslA* expression. Overexpression of *yfjC* resulted in a mucoid colony morphology and only slightly reduced *bslA* expression. These results indicate that YfjB and YfjC together inhibit DegSU activity.

The Δ*yfjD* mutant formed less-wrinkled colonies but had little or no effect on *bslA* expression in 2×SGG medium (Fig 7A, B). The less-wrinkled colony phenotype of the Δ*yfjD* mutant was more pronounced in MSgg medium (Fig S7), in which the Δ*yfjD* mutant had slightly decreased *bslA* expression. Since *yfjE* is a *yfjD* paralog, the Δ*yfjD*-*yfjE* (Δ*yfjDE*) double mutant was expected to have a stronger phenotype than the Δ*yfjD* mutant. However, the Δ*yfjDE* mutant had a phenotype similar to that of the Δ*yfjD* mutant in MSgg. The colony phenotype of the Δ*yfjDE* mutant was complemented by ectopic expression of *yfjD* (Fig S7), indicating that the Δ*yfjE* mutation does not alter colony morphology. Furthermore, Δ*yfjD* was suppressed by Δ*yfjC* because the Δ*yfjCDE* triple mutant formed WT-like wrinkled colonies. Thus, the colony phenotype of the Δ*yfjD* mutant is likely caused by the action of YfjBC.

### The *yfjA* operon encode a toxin–antitoxin system

YfjB contains transmembrane glycine zipper motifs that are found in pore-forming bacteriocins (Ali *et al*, 2023; Kim *et al*, 2005). I therefore hypothesized that the *yfjA* operon encodes a toxin− antitoxin system that mediates intercellular competition. To test this hypothesis, I carried out competition assays between WT and Δ*yfjA-*op strains. Cultures of the two strains were equally mixed based on OD_600_ values, and the mixture was grown on 2×SGG solid medium for 48 h. To distinguish the two competing strains, one constitutively expressed a *gfp* reporter, allowing quantification of the initial mixtures and the resulting colonies by flow cytometry. The *gfp* reporter did not affect competition because population changes were not observed in mixed colonies of WT+WT(*gfp*) or Δ*yfjA-*op+Δ*yfjA-*op(*gfp*) (Fig 8A). These assays demonstrated that the WT strain outcompeted the Δ*yfjA-*op mutant, with the percentage of the Δ*yfjA-*op mutant in WT+Δ*yfjA-*op mixed colonies decreasing to around 20% after 48 h (n=3). A decrease in the Δ*yfjA-*op mutant in WT+Δ*yfjA-*op colonies was also detected by fluorescence microscopy (Fig 8A). By contrast, competition between WT and Δ*yfjA-*op strains was not observed in MSgg solid medium (Fig 8A), indicating that, as observed in DegSU inhibition, high expression of the *yfjA* operon is required for competition.

**Figure 8.**
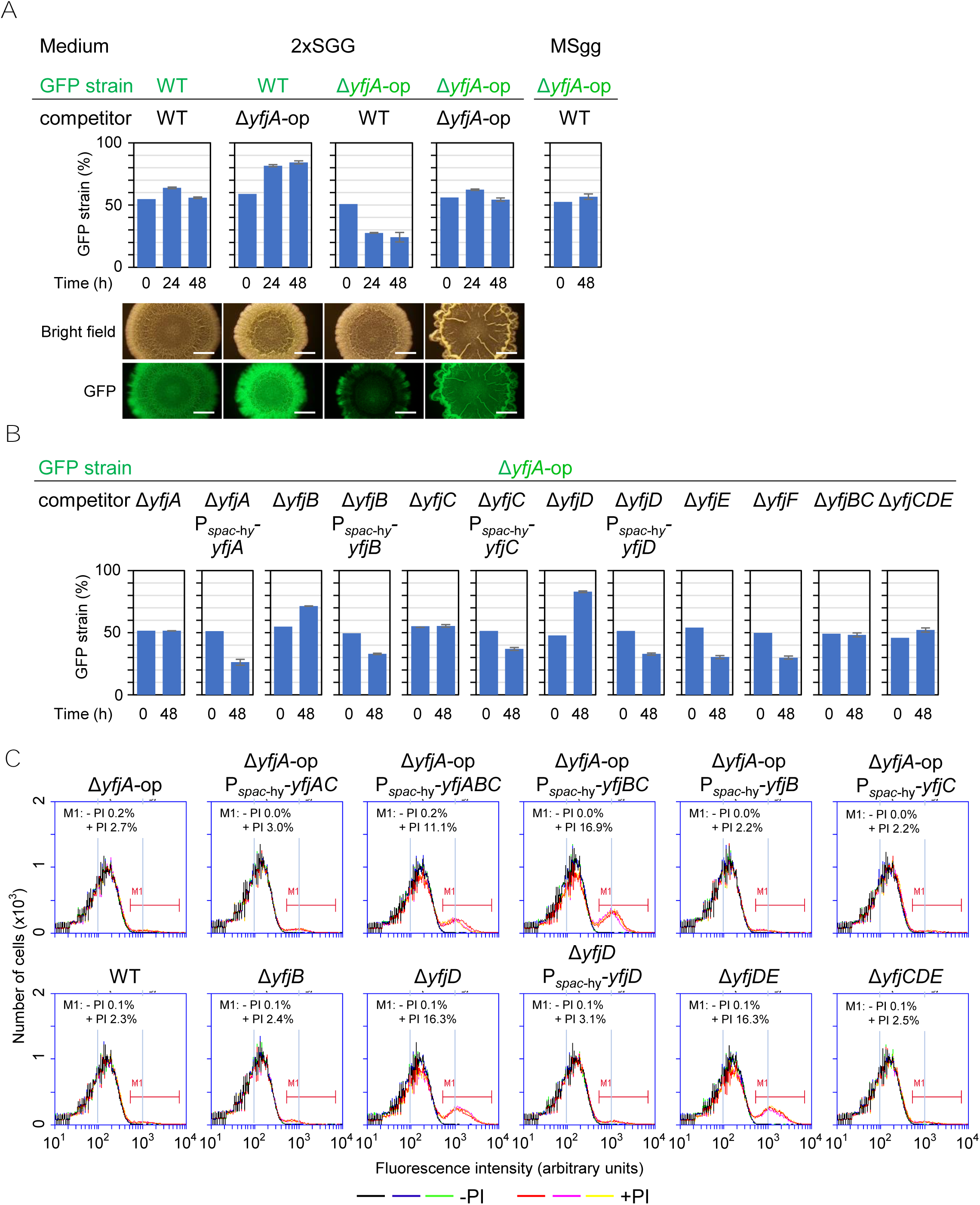
The *yfjA* operon mediates intercellular competition. A. Competition between WT and Δ*yfjA-*op strains. WT and Δ*yfjA-*op strains were grown to an OD_600_ of 0.7–0.8. One of the two strains was labeled with a constitutively expressed *gfp* reporter (GFP strains). Two cultures diluted to OD_600_ of 0.5 were mixed at a 1:1 ratio. Exact proportions of the GFP strains in the mixtures were determined by flow cytometry (time 0). The mixtures were spotted onto 2×SGG or MSgg medium and grown at 30°C. The proportion of GFP reporter strains within colonies at each time point was determined by flow cytometry using three independent colonies. Percentages are shown as mean values ± standard deviation (*n*=3). Resulting colonies were also analyzed by fluorescent microscopy. Brightfield and GFP images of colonies are shown below the graphs. Scale bar 2 mm. B. Identification of genes involved in attack on the Δ*yfjA-*op mutant. Competition assays of indicated strains were carried out as described in the panel A. Co-culture colonies including complemented strains were grown in 2×SGG supplemented with 10 µM IPTG. C. The YFJ toxin produces dead cell-containing colonies. Cells of 24 h-old colonies grown in 2×SGG were stained with PI and analyzed by flow cytometry. Average percentages (n=3) of PI-stained cells (M1) are indicated within plots. Plots without PI staining are shown as negative controls. Strains harboring the P*_spac_*_-hy_ construct were grown in 2×SGG supplemented with 100 µM IPTG.

To identify toxin and antitoxin genes, I co-cultured the Δ*yfjA-*op mutant with disruption mutants of individual *yfjA* operon genes. Δ*yfjA* and Δ*yfjC* mutants lost the ability to attack the Δ*yfjA-*op mutant (Fig 8B). The proportion of the Δ*yfjA-*op mutant was unchanged in co-culture colonies of Δ*yfjA-*op+Δ*yfjA* or Δ*yfjA-*op+Δ*yfjC*. By contrast, Δ*yfjB* and Δ*yfjD* mutants were outcompeted by the Δ*yfjA-*op mutant. Complementation of *yfjA*, *yfjB*, *yfjC*, and *yfjD* restored the inhibitory capacity of the corresponding mutants. *yfjE* and *yfjF* were not involved in competition because Δ*yfjE* and Δ*yfjF* mutants behaved like the WT strain in this assay.

Since *yfjA* and *yfjC* were required to outcompete the Δ*yfjA-*op mutant, I examined whether overexpression of *yfjAC* exerted toxicity in the Δ*yfjA-*op background. To this end, cells derived from 24 h-old colonies were stained with the living cell-impermeable fluorescent dye propidium iodide (PI), and the percentage of PI-stained dead cells were determined by flow cytometry. Contrary to expectations, overexpression of *yfjAC* did not promote cell death, with colonies consisting of 3% dead cells, similar to colonies of the parental Δ*yfjA-*op mutant (Fig 8C). Instead, overexpression of *yfjABC* promoted death in a subpopulation of cells with colonies containing >10% dead cells. Likewise, overexpression of *yfjBC* promoted cell death. However, overexpression of *yfjB* or *yfjC* alone did not promote cell death. These results indicate that YfjB and YfjC together exert toxicity in Δ*yfjA-*op mutant cells. I named the YfjBC-derived toxin YFJ. Since *B. subtilis* cells grown in liquid agitated culture take up PI (Kirchhoff & Cypionka, 2017), I could not determine whether overexpression of *yfjBC* causes immediate PI permeation. The observation that *yfjA* was required for intercellular competition but not toxicity indicates that the WXG100 family protein YfjA is likely required for YFJ toxin export.

Since the less-wrinkled colony phenotype of the Δ*yfjD* mutant was suppressed by the Δ*yfjC* mutation (see Fig S7), I speculated that the vulnerability of the Δ*yfjD* mutant in the competition assay is caused by susceptibility to self YFJ toxin. Consistent with this idea, Δ*yfjD* mutant colonies contained >10 % dead cells, and this phenotype was ablated in the *yfjD*-complemented strain (Fig 8C). Moreover, the Δ*yfjC* mutation suppressed cell death and interbacterial competition susceptibility phenotypes of the Δ*yfjD* mutant (see Δ*yfjCDE* in Fig 8B, C). These results indicate that YfjD functions as an antitoxin against the YFJ toxin. Although the Δ*yfjB* mutant did not exhibit significant cell death, the vulnerability of the Δ*yfjB* mutant in the competition assay was suppressed by the Δ*yfjC* mutation (see Δ*yfjBC* in Fig 8B). I speculate that the chaperone homolog YfjC may interact with some cellular proteins in the absence of YfjB, causing weak adverse effects on cell growth.

### T7SS is required for YFJ toxin-mediated competition

Sequence features of YfjA and YfjC suggested that the YFJ toxin is exported by a T7SS, the components of which are encoded by the *yukE* operon. This idea is supported by the observation that deletion of the entire *yukE* operon (Δ*yukE*-op) abolished competition between WT and Δ*yfjA*-op strains (Fig 9A).

**Figure 9.**
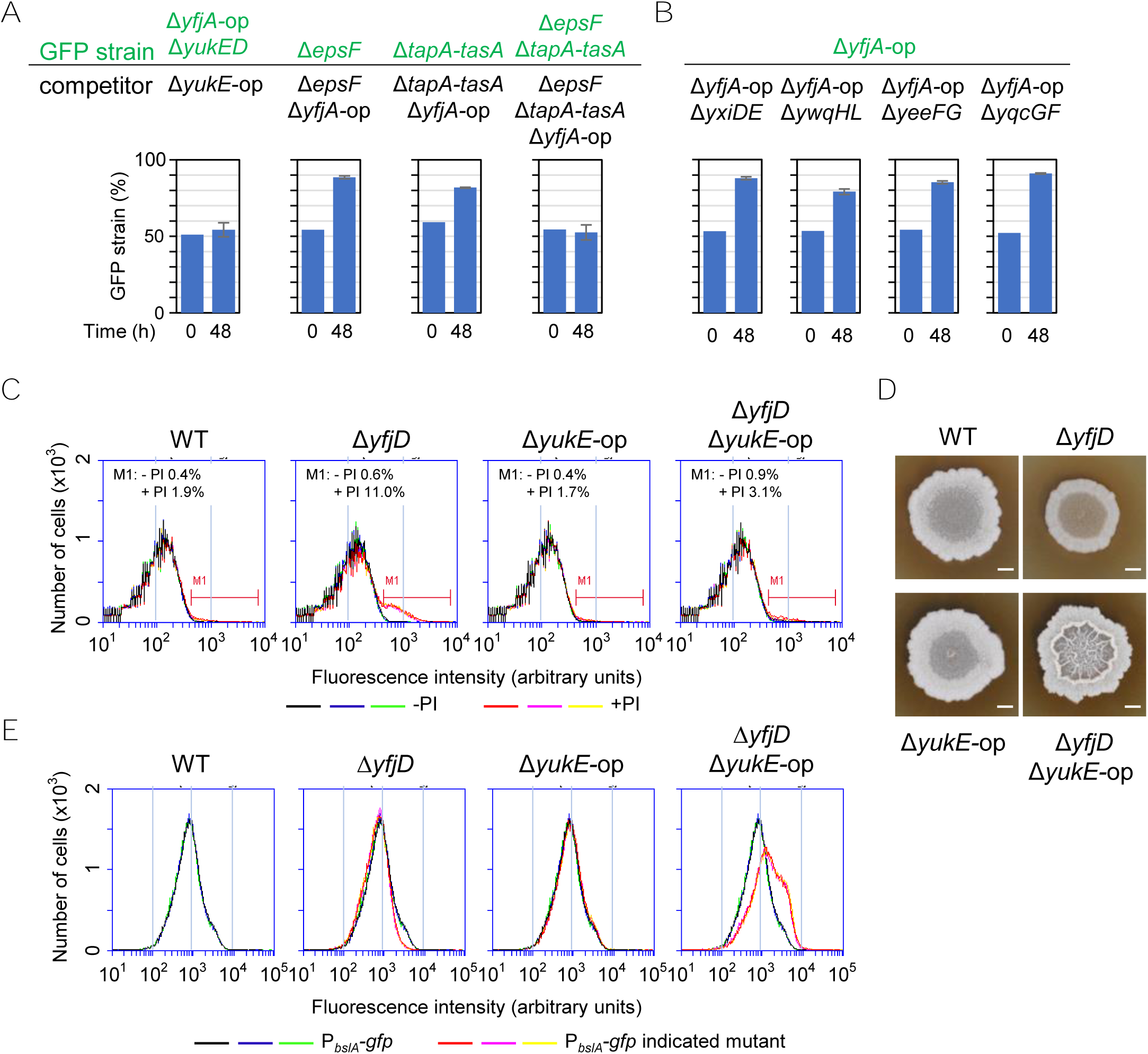
The YFJ toxin is exported by the T7SS. A. Competition assays. Competition assays of indicated strains were carried out as described in Fig 8A. B. The Δ*yfjA*-op mutation does not inhibit LXG toxin-mediated competition. C. The Δ*yukE*-op mutation suppresses cell death in the Δ*yfjD* mutation. Cells of 24 h-old colonies grown in 2×SGG were stained with PI and analyzed by flow cytometry. Average percentages (n=3) of PI-stained cells (M1) are indicated within plots. Plots without PI staining are shown as negative controls. D. The colony morphology of the Δ*yfjD* Δ*yukE*-op mutant. Strains were grown in 2×SGG medium for 48 h. Scale bar, 2 mm. E. Expression of P*_bslA_-gfp* at 24 h in the Δ*yfjD* Δ*yukE*-op mutant. *bslA* expression in WT is shown as a reference.

The T7SS and its known substrates, LXG toxins, are induced by DegSU and function specifically in biofilms (Kobayashi, 2021). I therefore determined whether the *yfjA* operon system also functions specifically in biofilms. To this end, I carried out competition assays between WT and Δ*yfjA-*op in Δ*epsF* and/or Δ*tapA-tasA* mutation backgrounds, which are deficient in major EPS components, exopolysaccharides and TasA amyloid fibers, respectively. Although neither the Δ*epsF* mutation nor the Δ*tapA-tasA* mutation alone affected competition between WT and Δ*yfjA-*op strains, the Δ*epsF* Δ*tapA-tasA* combination abolished competition (Fig 9A). The Δ*epsF* Δ*tapA-tasA* combination did not prevent *yfjA* operon expression (Fig S8). Thus, delivery of the YFJ toxin depends on exopolysaccharides and TasA amyloid fibers. These results indicate that the YFJ toxin system adapts to the biofilm environment and functions specifically in biofilms similar to LXG toxin systems.

I next examined whether the Δ*yfjA-op* mutation prevents LXG toxin-dependent competition. Competition assays showed that Δ*yfjA-op* mutations did not prevent any of the four LXG toxins from attacking the deletion mutants of LXG toxin–antitoxin operons (Fig 9B), indicating that T7SS-dependent YFJ and LXG toxins are functionally separable.

I isolated transposon insertions that suppressed cell death in the Δ*yfjD* mutant. One transposon was inserted in the middle of *yukB*, the fourth gene of the *yukE* operon. Indeed, the Δ*yukE-*op mutation suppressed cell death in the Δ*yfjD* mutant (Fig 9C). These results indicate that the YFJ toxin loses its toxicity in the Δ*yfjD* Δ*yukE-*op mutant. Since some bacteriocins require membrane receptors for their action (Diep *et al*, 2007), YfjD and *yukE* operon proteins may function as receptors for the YFJ toxin. However, since the Δ*yukE-*op mutation prevented YFJ toxin export, these proteins may function as receptors for intracellular YFJ toxin, and thus it is unclear whether these proteins function as receptors for extracellular YFJ toxin. The Δ*yfjD* Δ*yukE-*op mutant formed macroscopically wrinkled colonies and exhibited increased expression of *bslA* (Fig 9D, E), like the Δ*yfjA*-op mutant. Thus, the Δ*yfjD* Δ*yukE-*op mutation impairs both YFJ toxin toxicity and inhibitory activity against DegSU. Considering that *yfjB* and *yfjC* are responsible for both cellular toxicity and the inhibition of DegSU, these results indicate that the YFJ toxin causes both cellular toxicity and DegSU inhibition.

### The YFJ toxin system functions as an intercellular signal for DegSU

I hypothesized that the secreted YFJ toxin might control DegSU activity as an intercellular signal. To test this hypothesis, I co-cultured the Δ*yfjA-*op mutant harboring the P*_bslA_-gfp* reporter (referred to as Δ*yfjA-*op P*_bslA_-gfp*) with the WT strain or the Δ*yfjA-*op mutant at various initial mixing ratios before analyzing the fluorescence profiles of the resulting colonies by flow cytometry. Since the WT strain dominated the Δ*yfjA-*op mutant in co-culture colonies, initial mixing rates were modified. The co-culture colonies exhibited two fluorescence signals, the GFP fluorescence from Δ*yfjA-*op P*_bslA_-gfp* cells and the autofluorescence of the WT or Δ*yfjA-*op cells (Fig 10). The intensity of GFP fluorescence in Δ*yfjA-*op P*_bslA_-gfp*+Δ*yfjA-*op co-culture colonies was comparable to that in Δ*yfjA-*op P*_bslA_-gfp* mono-culture colonies (initial ratio of 10:0), regardless of the initial mixing ratio. By contrast, the GFP fluorescence intensity in Δ*yfjA-*op P*_bslA_-gfp*+WT co-culture colonies became lower as the initial mixing ratio of the WT strain was increased. Thus, the YFJ toxin produced by WT cells reduced DegSU activity in Δ*yfjA-*op mutant cells. These results indicate that the YFJ toxin can function as an intercellular signal for DegSU.

**Figure 10.**
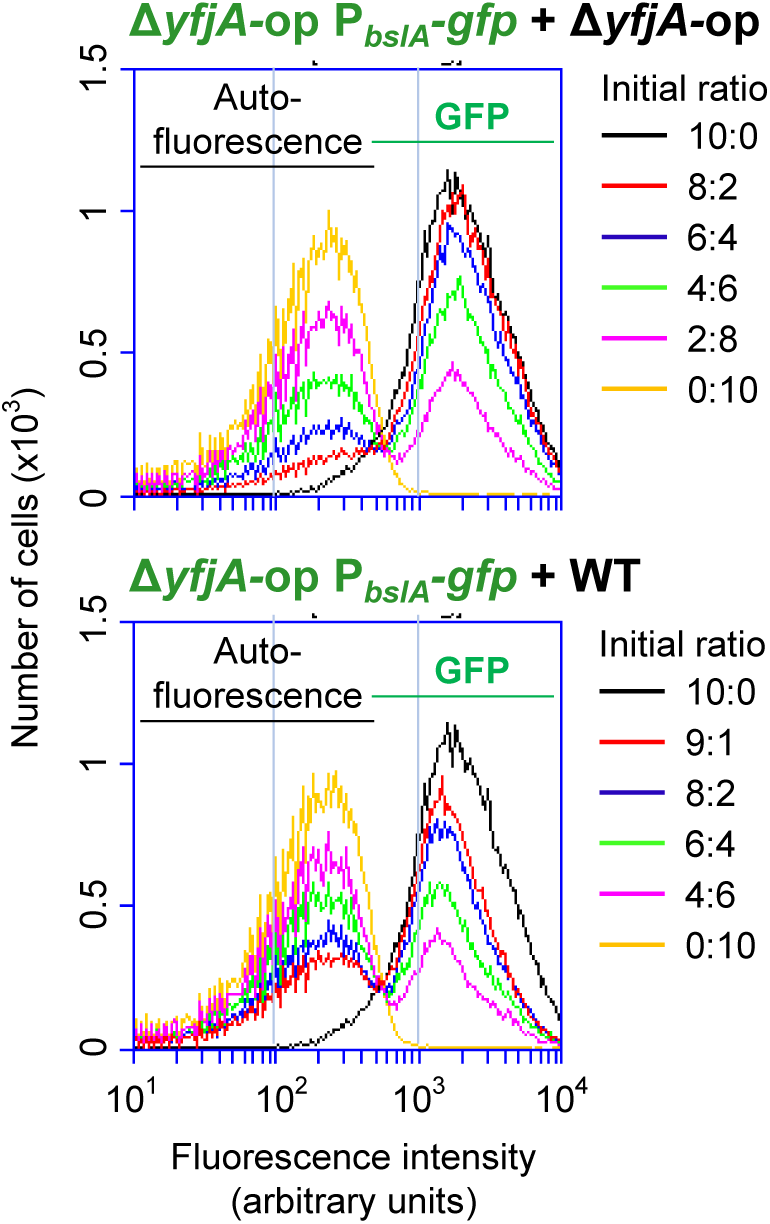
The YFJ toxin can intercellularly inhibit DegSU activity. The Δ*yfjA*-op mutant harboring the P*_bslA_-gfp* reporter was mixed with the Δ*yfjA*-op mutant (Top) or the WT strain (Bottom) at indicated initial ratios, and the two strains were co-cultured for 24 h. Fluorescence profiles of resulting colonies were analyzed by flow cytometry. Peaks of autofluorescence and GFP are indicated within plots. 10:0 and 1:0 plots are common between the two panels. Three colonies were analyzed for each initial ratio, and represented plots are shown.

## Discussion

In this study, I attempted to identify an intercellular signaling mechanism adapted to the biofilm environment. My results demonstrated that the *yfjA* operon encodes a T7SS effector toxin, named the YFJ toxin, which acts as both a secreted toxin and an intercellular signal for the DegSU two-component system in colony biofilms. Considering that T7SS-dependent LXG toxins are widespread as biofilm-specific weapons in *B. subtilis* (Kobayashi, 2021), I propose that contact-dependent toxins and signaling can function as biofilm-adapted mechanisms.

The *yfjA* operon encodes the YFJ effector toxin and its related proteins. Ectopic expression of YfjBC caused permeation of the membrane-impermeable dye PI in >10% of cells in colony biofilms. Since PI permeation is an indicator of membrane damage, this phenotype is consistent with observations that YfjB contains transmembrane glycine zipper motifs commonly found in pore-forming toxins. In pore-forming toxins, glycine zipper motif domains permeate the membrane of target cells, forming pores (Kim *et al*, 2004; Kim *et al*, 2005) or facilitate the translocation of toxin domains into cells (Ali *et al*, 2023). As YfjB seems to have no additional toxin domain, presence of these motifs suggest that YfjB may function as a pore-forming toxin. However, YfjB exhibited toxicity only in the presence of YfjC. Since YfjC is structurally similar to the EspG chaperone of the mycobacterial T7SS, it seems unlikely that YfjC is directly involved in toxicity. YfjC may be required for other functions, such as the stabilization and multimerization of YfjB.

YFJ toxin-mediated competition was observed in nutrient-rich 2×SGG medium but not in minimal MSgg medium. Likewise, the YFJ toxin inhibited DegSU activity only in 2×SGG medium. Since the *yfjA* operon was more strongly expressed in 2×SGG medium than in MSgg medium, I speculate that YFJ toxin action is dependent on high levels of toxin production. The glycine zipper toxin CdzC of *Caulobacter crescents* forms large aggregates on the surface of producer cells to attack recipient cells (García-Bayona *et al*, 2017). Likewise, the YFJ toxin may also need to accumulate on the cell surface to attack recipient cells, and expression levels of the *yfjA* operon in MSgg medium may be insufficient for such accumulation.

The T7SS was required for YFJ toxin-mediated intercellular competition. YfjA and YfjC are homologous to known T7SS chaperones. These results suggest that the YFJ toxin is exported by the T7SS with the help of YfjA and YfjC. The T7SS was shown to deliver effector toxins to recipients in a cell–cell contact-dependent manner in several Gram-positive bacteria (Bowran & Palmer, 2021; Cao *et al*, 2016; Chatterjee *et al*, 2021; Ulhuq *et al*, 2020). In *B. subtilis*, the T7SS delivers LXG toxins to recipient cells and is dependent on exopolysaccharides and TasA amyloid fibers (Kobayashi, 2021). Since these EPS components hold cell chains close together and thereby align cells in biofilms (Branda *et al*, 2006; Branda *et al*, 2001; Kobayashi, 2007a; Romero *et al*, 2010), these EPSs are thought to facilitate the delivery of LXG toxins to recipient cells by stabilizing cell–cell contact (Kobayashi, 2021). YFJ toxin-mediate intercellular competition also depended on exopolysaccharides and TasA amyloid fibers, suggesting that the YFJ toxin is also exported by T7SS in a cell–cell contact-dependent manner. I have previously showed that LXG toxins are numerous and highly diverse in *B. subtilis* strains and are used as weapons in biofilms. The finding of another class of T7SS substrates indicates that cell–cell contact-dependent delivery systems may have advantages over general secretion systems in biofilms. In biofilms, each bacterial strain often forms its own cluster, with progeny cells remaining close parent cells (Nadell *et al*, 2016). Therefore, in biofilms, cell–cell contact-dependent delivery systems may be more effective through delivery of toxins or signal molecules to neighboring cells, rather than to the entire community.

The YFJ toxin suppressed non-immune cells lacking the antitoxin YfjD. However, the YFJ toxin had only weak toxicity and did not prevent colony formation even upon overexpression. The YFJ toxin is a predicted bacteriocin, which usually has a narrow range of activity directed toward related strains. Bacteriocins exhibit polymorphism and are unevenly distributed among related strains, contributing to self-recognition (Wall, 2016). However, the *yfjA* operon is completely conserved among the 13 *B. subtilis* strains tested, and *yfjA* operon proteins showed little sequence variation among the 13 strains. Since the *yfjA* operon was uniformly expressed in 2×SGG and MSgg media, the YFJ toxin does not appear to function as a cannibalism toxin (González-Pastor *et al*, 2003). The YFJ toxin is therefore unlikely to be toxic in biofilms. Instead, the YFJ toxin inhibited DegSU not only in non-immune cells but also in WT cells expressing the antitoxin YfjD. I therefore propose that the YFJ toxin primarily functions as an intercellular signal. The high conservation of the *yfjA* operon suggests that the YFJ toxin may serve species-specific, rather than strain-specific signaling functions. The inhibition mechanism of DegSU by the YFJ toxin is currently unknown. Expression of DegSU-regulated genes are induced by NaCl stress, methyl salicylate, and the inhibition of flagellar rotation (Cairns *et al*, 2013; Chan *et al*, 2014; Kobayashi, 2015; Steil *et al*, 2003), all of which affect the cell membrane or membrane-related functions. I therefore speculate that the YFJ toxin may cause membrane changes even in the presence of the YfjD antitoxin protein, thus preventing DegSU activation.

The YFJ toxin system modulated DegSU but was not solely responsible for its activity Modulation of DegSU activity not only changes expression of DegSU-regulated genes but also changes growth phenotype. During growth and development, DegU∼P levels gradually increase and incrementally activates genes involved in motility, biofilm formation, and sporulation (Kobayashi, 2007b; Marlow *et al*, 2014b; Verhamme *et al*, 2007). Promoters of genes involved in these functions have different affinities for DegU∼P, which determines the temporal order of gene expression during growth and development (Kobayashi, 2007b). Therefore, the modulation of DegSU activity is involved in cell fate decisions, and an increase in DegSU activity in biofilms not only increases expression of DegSU-regulated genes but also promotes the transition from biofilm growth to the sporulation/dormant mode.

DegSU strongly activated expression of DegSU-regulated genes including the *yfjA* operon in nutrient-rich biofilm medium 2×SGG (Kobayashi, 2021), promoting the inhibitory activity of the YFJ toxin system on DegSU. Expression of *degU* was reported to be enhanced by the catabolite control protein CcpA in glucose-rich medium (Ishii *et al*, 2013). Since 2×SGG medium contains multiple carbon sources (1% glycerol, 0.1% glucose, and carbohydrates from nutrient broth), this mechanism of regulation likely contributes to the strong expression of DegSU-regulated genes in 2×SGG medium. DegSU is also required for efficient glucose utilization, which promotes the release of CcpA-dependent catabolite repression, leading to glucose-resistant sporulation (Kunst *et al*, 1974; Tanaka *et al*, 2015). These observations are consistent with the findings that increased DegSU activity in the Δ*yfjA*-op mutant hastened and promoted sporulation in carbon-rich 2×SGG medium. I propose that the *yfjA* operon is strongly induced under carbon-rich conditions, regulating DegSU activity in biofilms, thereby prolonging the biofilm growth period. Delaying sporulation and continued growth may be advantageous for *B. subtilis* to proliferate in competitive biofilms under nutrient-rich environments.

Intercellular signaling via specialized secretion systems has been reported previously. The type VI secretion system in *Yersinia pseudotuberculosis* delivers the effector toxin CccR to recipient cells, where CccR acts both as a toxin and as a transcriptional regulator (Wang *et al*, 2022). The contact-dependent growth inhibition toxin BcpA in *Burkholderia thailandensis* induces changes in transcription of hundreds of genes in recipient cells (Garcia *et al*, 2016). However, unlike these toxins, the YFJ toxin system functioned specifically in biofilms. Several possible scenarios pertain to the function of the YFJ toxin system in biofilms. Firstly, when a *B. subtilis* cell forms a large cluster with its siblings in biofilms, the cell may be attacked by a large quantity of the YFJ toxin from surrounding cells, leading to inhibition of DegSU activity, prolonging the biofilm growth period. Instead, when *B. subtilis* cells live as a minority surrounded by competitors in biofilms, the YFJ toxin is not delivered to the cell sufficiently to inhibit DegSU activity, and thus the cell initiates sporulation early even under nutrient-rich conditions. This complex regulatory system is likely a survival strategy allowing *B. subtilis* to thrive in adverse environments.

## Materials and methods

### Bacterial strains and media

*B. subtilis* strain NCIB 3610 and its derivatives used in this study are listed in Appendix Table S1. The construction of *B. subtilis* mutants is described in the Appendix. Primers used for strain construction are listed in Appendix Table S2. *B. subtilis* strains were grown in 2×SGG [16 g/L nutrient broth (BD Difco, Franklin Lakes, NJ, USA), 1% glycerol, 0.1% glucose, 0.2% (w/v) KCl, 2 mM MgSO_4_·7H_2_O, 1 mM Ca(NO_3_)_2_, 0.1 mM MnCl_2_, 1 µM FeSO_4_] (Kobayashi, 2007a), MSgg [5 mM potassium phosphate (pH 7), 100 mM MOPS (pH 7), 2 mM MgCl_2_, 700 μM CaCl_2_, 50 μM MnCl_2_, 50 μM FeCl_3_, 1 μM ZnCl_2_, 2 μM thiamine, 0.5% (w/v) glycerol, 0.5% (w/v) glutamate, 50 μg/mL tryptophan] (Branda *et al*, 2001), or LB [LB Lennox; 10 g/L bacto tryptone (BD Difco), 5 g/L yeast extract (BD Difco), 5 g/L NaCl]. *E. coli* strains JM105 and JM109 were used for the construction and maintenance of plasmids.

### Bioinformatic analysis

A Kyte-Doolittle hydropathy plot of YfjB was constructed using Expasy (https://web.expasy.org/protscale/). The transmembrane regions of YfjB were analyzed using PSORT (https://psort.hgc.jp/), PRED-TMR (http://athina.biol.uoa.gr/PRED-TMR/), SPLIT (http://split.pmfst.hr/split/4/), and HMMTOP (http://www.enzim.hu/hmmtop/index.php). Multiple sequence alignments were performed using CLUSTALW (https://www.genome.jp/tools-bin/clustalw) with default settings. Gene organization was compared using SyntTax (https://archaea.i2bc.paris-saclay.fr/syntax/).

### Growth conditions for biofilm formation

Overnight cultures (150 μL) of *B. subtilis* strains at 28°C in LB were added to 5 mL LB, and the strains were grown with shaking at 37°C until the OD_600_ reached 0.5–0.8. The cultures were then diluted to an OD_600_ of 0.05 with LB, and 2 µL of the dilutions were spotted onto solid medium. The inoculated plates were then incubated at 30°C. For pellicle formation, 100 µL of each dilution was added to 10 mL 2×SGG medium in a well of a 6-well dish, and the dishes were incubated at 30°C.

### Expression of *gfp* reporters

Colonies were grown as described above. One colony was scraped with an inoculation loop and suspended in 600 μL PBS (81 mM Na_2_HPO_4_, 26.8 mM KCl, 14.7 mM KH_2_PO_4_, 1.37 mM NaCl,). After cell aggregates in the suspension were dispersed by mild sonication, the cells were pelleted by centrifugation at 5,800 × g for 1 min. The cells were then fixed with 4% paraformaldehyde for 7 min and washed with PBS as previously described (Vlamakis *et al*, 2008). Single-cell fluorescence was measured using an Accuri C6 flow cytometer (BD Biosciences, Franklin Lakes, NJ, USA). Fifty thousand events were recorded per sample. The threshold for FSC-H was set at 20,000 to remove signals from cell debris and contaminants. Three independent colonies were analyzed for each test.

### GFP fluorescence in SDS-PAGE

Four colonies grown on 2×SGG for 24 h were suspended in 700 μL PBS supplemented with 10 mM EDTA (pH 8.0) and 2 mM phenylmethylsulfonyl fluoride (PMSF). After mild sonication to disperse cell aggregates, samples were diluted to an OD_600_ of 8.6. Five hundred microliters of diluted samples were mixed with 5 μL of 10% SDS and sonicated to lyse the cells. After centrifugation at 17,400 × g for 5 min, 8 μL supernatant was separated by SDS-PAGE (SuperSep Ace 10–20%, FUJIFILM Wako Pure Chemical, Osaka, Japan) without pre-heat treatment. After electrophoresis, GFP fluorescence images were captured immediately with FUSION FX (Vilber, Marne-la-Vallée, France). The gels were then stained with Coomassie Brilliant Blue R250, and the stained images were captured using FUSION FX. The gel images were quantitatively analyzed using Evolution Capt (Vilber).

### Microscopy

Expression of *gfp* reporters in *B. subtilis* colonies was analyzed using an SZX7 stereomicroscope (Olympus, Tokyo, Japan) equipped with an AdvanCam-E3Rs digital color camera (Advan Vision, Tokyo, Japan). Images were obtained using AdvanView (Advan Vision), and the file size was reduced for figures using Photoshop Elements (Adobe, San Jose, CA, USA). The experiments were performed at least twice, with similar results.

### Co-culture assay

*B. subtilis* strains were grown to late-exponential phase as described above. Two culture dilutions (500 µL, OD_600_ of 0.5) were mixed well by vortexing. Two microliters of each mixture was spotted on 2×SGG solid medium, and the remaining volume was used to determine the proportions of the two strains at time 0 by flow cytometry. The inoculated plates were incubated at 30°C. After 24 h or 48 h, colonies were harvested and fixed to determine the proportions of the two strains by flow cytometry. The proportions of the two strains in the mixed colonies were determined by averaging the data from three colonies.

### PI staining

A 24 h-old colony was suspended in 600 μL PBS, and cell aggregates in the suspension were dispersed by mild sonication. 98 μL suspension was then mixed with 2 μL of 50 μg/mL PI (FUJIFILM Wako Chemicals, Osaka, Japan). After 10 min of incubation, single-cell fluorescence was measured using an Accuri C6 flow cytometer, as described above.

### Transposon mutagenesis

The temperature-sensitive vector pMarA carrying Tn*YLB* (Le Breton *et al*, 2006) was introduced into the Δ*yfjD* mutant, which formed translucent colonies on LB solid medium. four independent colonies were cultured at 28°C for 18 h in LB supplemented with kanamycin. The cultures were diluted 10-fold and further cultured at 43°C for 2 h. This process was repeated once more. After centrifugation at 5,800 × g for 5 min, cells were suspended in LB 20% glycerol and stored at −80°C. The stocks were serially diluted and spread on LB solid medium supplemented with kanamycin, and WT-like colonies were screened at 37°C. To confirm that the transposon insertions were responsible for suppressor activity, chromosomal DNA from the suppressor mutants was used to transform the Δ*yfjD* mutant. To remove transposons inserted in the *yfjA* operon, the chromosomal DNA was used to transform the WT strain, and transposon insertions linked to the Δ*yfjD* mutation were removed. Transposon insertion sites were determined according to the procedure described by Le Breton *et al* (2006).

## Appendix

### Construction of *B. subtilis* strains

Since strain NCIB 3610 has low competence ability, mutant alleles were first introduced into the domesticated strain 168 and then transferred to strain NCIB 3610 via transformation with genomic DNA (4). Since strain 168 has several mutations that affect biofilm formation, the possibility of introducing these unwanted biofilm defects into strain NCIB 3610 was eliminated by examining the colony morphology of transformants.

### The P*_yfjA_*-*gfp* transcriptional fusion

The promoter region of *yfjA* was amplified by PCR with primers, *yfjA*-P-F2 and *yfjA*-P-R2. The PCR products were digested with *BamH*I and *Hin*dIII, and then ligated with the *BamH*I- and *Hin*dIII-digested plasmid pDCG3, which is an *amyE* integration vector containing promoterless *gfp* and *cat* genes. The resultant plasmids were used to transform strain 3610, generating the strain N1791 (*amyE*::P*_yfjA_-gfp*, *cat*).

### The *htpG*::P*_bslA_-gfp* transcriptional fusion

The promoter region of *bslA* was amplified by PCR with primers, *bslA-*P-F4 and *bslAA*-P-R4. The PCR products were digested with *BamH*I and *Hin*dIII, and then ligated with the *BamH*I- and *Hin*dIII-digested plasmid pHCG3, which is an *htpG* integration vector containing promoterless *gfp* and *cat* genes. The *cat* maker was replaced with the *tet* maker using pCM::TET (12)

### Deletion mutants

Deletion of the *yfjA* operon was carried out using an overlap extension PCR technique. A *cat* cassette was amplified from pCBB31 by PCR with pUC-R and pUC-F primers (Table S2). Upstream and downstream regions of the *yfjA* operon were amplified by PCR with primer pairs *yfjA*-F1–*yfjA*-R1 and *yfjE*-F2–*yfjE*-R2, respectively. The 5′ sequences of *yfjA*-R1 and *yfjE*-F2 were complementary to the sequences of pUC-R and pUC-F, respectively. To fuse the three PCR fragments, all three were mixed and used as template for a second round of PCR using primers *yfjA*-F1 and *yfjE*-R2. The resultant PCR products were used to transform strain 168, generating the strain 168 Δ*yfjA-*op::*cat*. The *cat* maker was then replaced to *spc* by transforming with plasmid pCM::SP (12). The Δ*yfjA-* op::*spc* mutation was then transferred to strain 3610, generating N1451 (Δ*yfjA-*op::*spc*).

Deletion mutants of Δ*yfjDE,* Δ*yfjCDE*, *and* Δ*epsF* were constructed via the same procedure using different sets, Δ*yfjDE, yfjD*-F1, *yfjD*-R1, *yfjE*-F2, and *yfjE*-R2; Δ*yfjCDE, yfjC-F6, yfjC*-R6, *yfjE*-F2-2, and *yfjE*-R2-2; Δ*epsF*, *epsF*-F1, *epsF*-R1, *epsF*-F2, and *epsF*-R2 (Table S2). The *cat* marker of Δ*yfjDE* was replaced to *erm* by transforming with plasmid pCM::EM (12).

### Δ*yfjA*, Δ*yfjB,* and Δ*yfjC* maker-less, in-frame deletion mutants

To delete *yfjA*, upstream and downstream regions of *yfjA* were amplified using primer sets, *yfjA*-D-F1–*yfjA*-D-R1 and *yfjA*-D-F2–*yfjA*-D-R2, respectively (Table S2). 5’ parts of *yfjA*-D-R1 and *yfjA*-D-F2 primers complementary to each other. Two PCR products were mixed and used as template for a second round of PCR using primers *yfjA*-D-F1 and *yfjA*-D-R2. The resultant PCR products were digested with *Eco*RI and *Bam*HI, and then ligated with the *BamH*I- and *Hin*dIII-digested plasmid pMAD (1). This plasmid was introduced into NCIB3610 by transformation. Insertion and excision of pMAD*yfjA* into the chromosome of NCIB3610 was performed as previously described (7). The deletion of *yfjA* was confirmed by PCR. The deletion mutants of *yfjB* and *yfjC* were constructed via the same procedure using different primer pairs, *yfjB*, *yfjB*-D-F1–*yfjB*-D-R1 and *yfjB*-D-F2–*yfjB*-D-R2; *yfjC*, *yfjC*-D-F1–*yfjC*-D-R1 and *yfjC*-D-F2–*yfjC*-D-R2 (Table S2).

### Δ*yfjD*::pMutin, Δ*yfjE*::pMutin, and Δ*yfjF*::pMutin mutants

pMutin vector insertion mutants of *yfjD*, *yfjE*, and *yfjF* genes were previously reported (8). The *erm* maker of pMutin was replaced to *tet* by transforming these mutants with plasmid pEM::TC (12), and then the resultant mutations were transferred to strain 3610.

### *P_spac_-degU* and *P_spac-hy_-yfjA* operon strains

5’ parts of *degU* and *yfjA* including the SD sequence were amplified by PCR with primer pairs, *degU*-F10–*degU*-R10 and *yfjA-F2*– *yfjA-R2*, respectively. PCR products were digested with *Hin*dIII and *Bam*HI and then ligated with the *BamH*I- and *Hin*dIII-digested plasmid pMutinNC (10) and pMutinT3-hy (4), respectively. Resultant plasmids were used to construct N2622 (*P_spac_-degU*) and N2613(*P_spac-hy_-yfjA* operon) strains.

### *amyE*::P*_spac-hy_-yfjA, amyE*::P*_spac-hy_-yfjB, amyE*::P*_spac-hy_-yfjC, amyE*::P*_spac-hy_-yfjD, amyE*::P*_spac-hy_-yfjABC, and amyE*::P*_spac-hy_-yfjBC*

*yfjA* was amplified by PCR using chromosomal DNA of NCIB3610 and primers, *yfjC*-F2-3 and *yfjA*-R3-2. 5’ sequences of *yfjA*-F2-2 and *yfjA*-R3-2 are complementary to the upstream sequence of the *Hin*dIII site on pDLT3-hy (9) and the downstream sequence of the *Bam*HI site on pDLT3-hy, respectively. The PCR products were mixed with *Hin*dIII and *Bam*HI-digested pDLT3-hy, and two DNA fragments were ligated at 50°C for 2 h using the Gibson assembly master mix (New England Biolabs, Massachusetts, USA). The ligation mixtures were then used to transform the strain 168. Among Cm^r^ transformants, the *yfjA*-expression strain was identified by PCR. The expression construct in the resultant strains was then transferred to the strain NCIB 3610. The *yfjB*-, *yfjC*-, *yfjD-*, *yfjABC-, yfjBC-*expression strains were constructed via the same procedure using different primer pairs, *yfjB*, *yfjB*-F3 and *yfjB*-R3-2; *yfjC*, *yfjC*-F3-2 and *yfjC*-R3-2; *yfjD*, *yfjD*-F4 and *yfjD*-R4; *yfjABC*, *yfjA*-F2-2 and *yfjC*-R3-2. For the construction of the *yfjAC*-expression strain, DNA was amplified via PCR using the chromosomal DNA of the Δ*yfjB* mutant and primers, *yfjA*-F2-2 and *yfjC*-R3-2. These constructs were transferred to deletion mutants of *yfjA* operon genes.

---

**Table S1.**
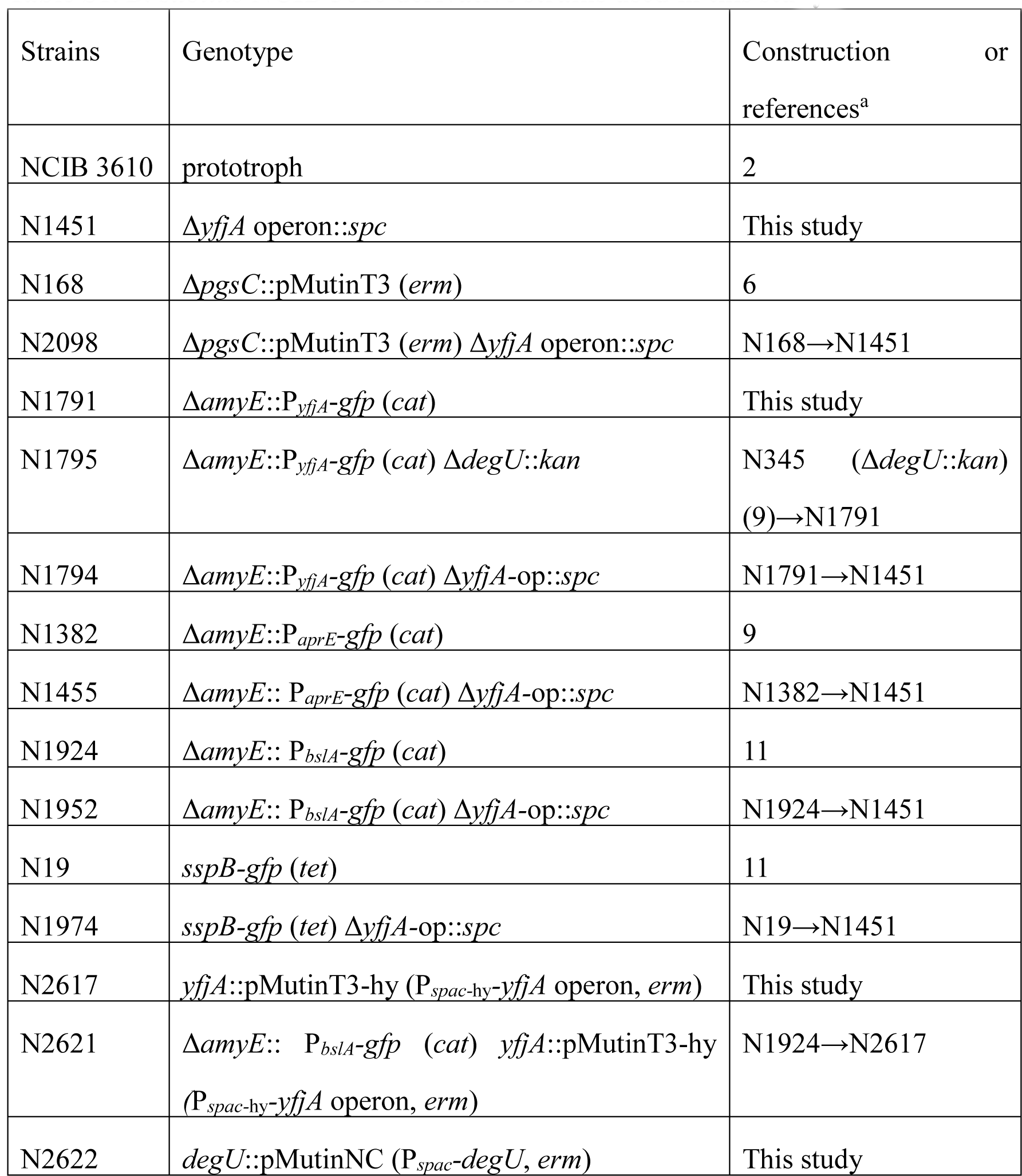
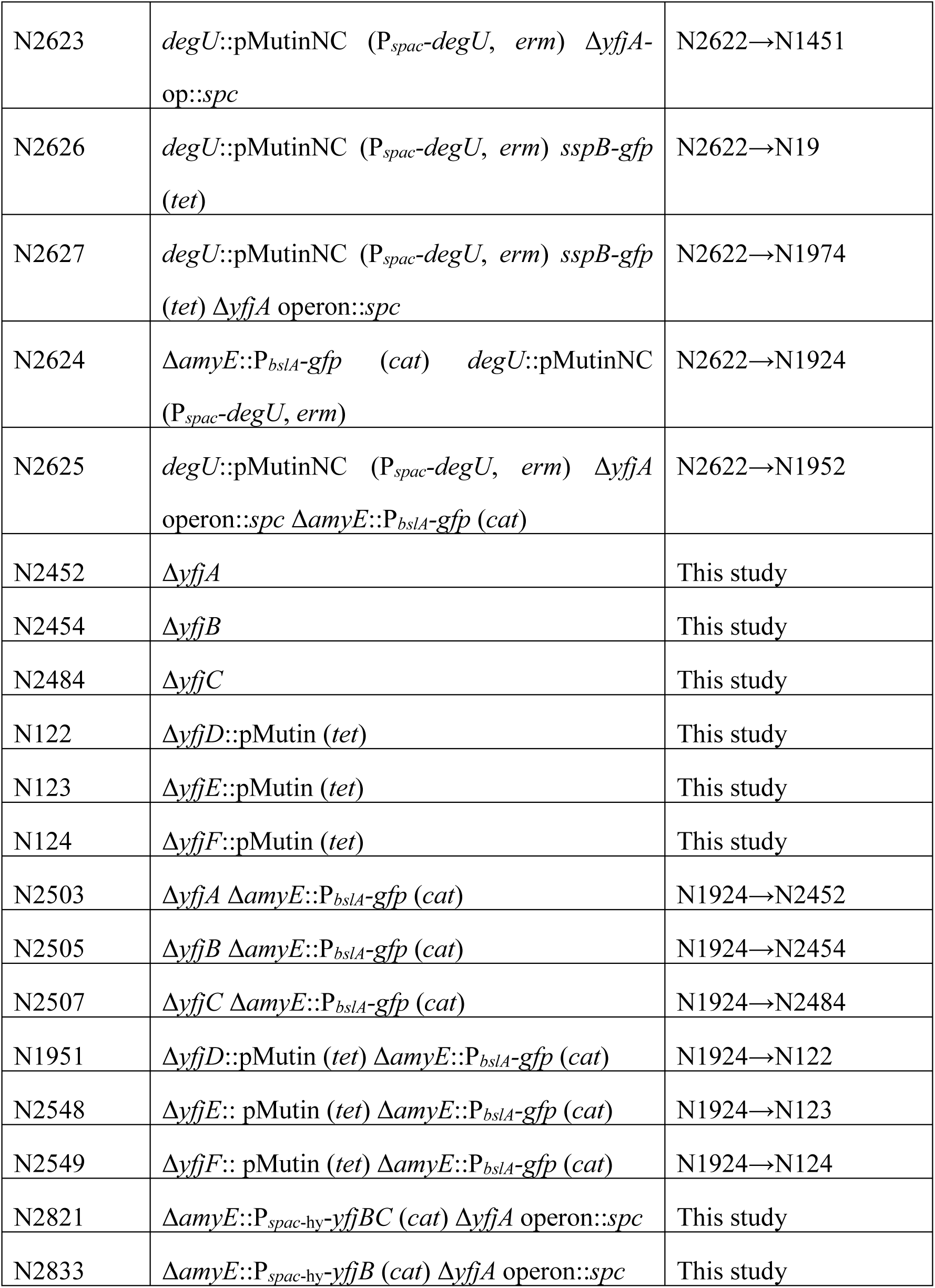
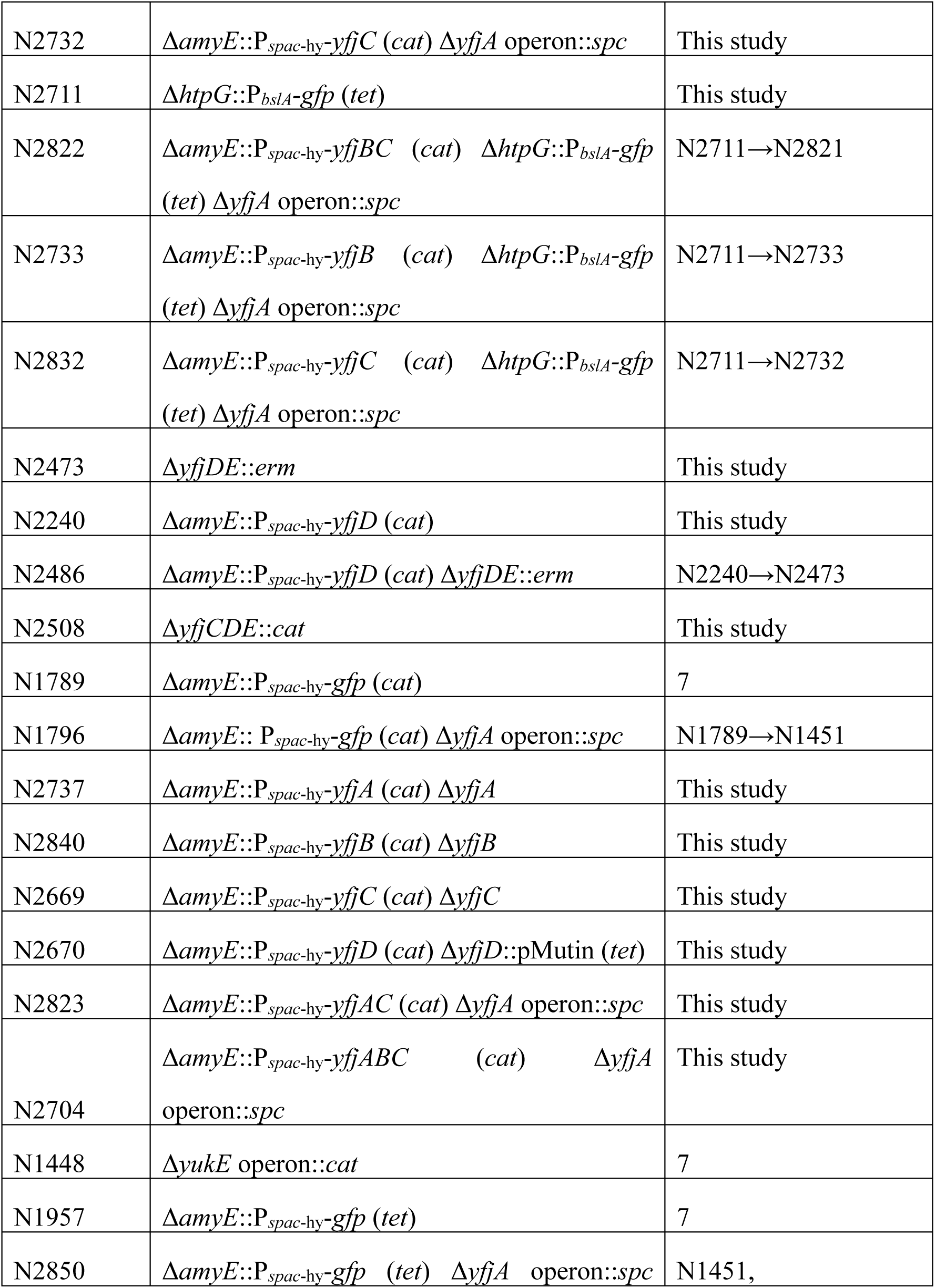
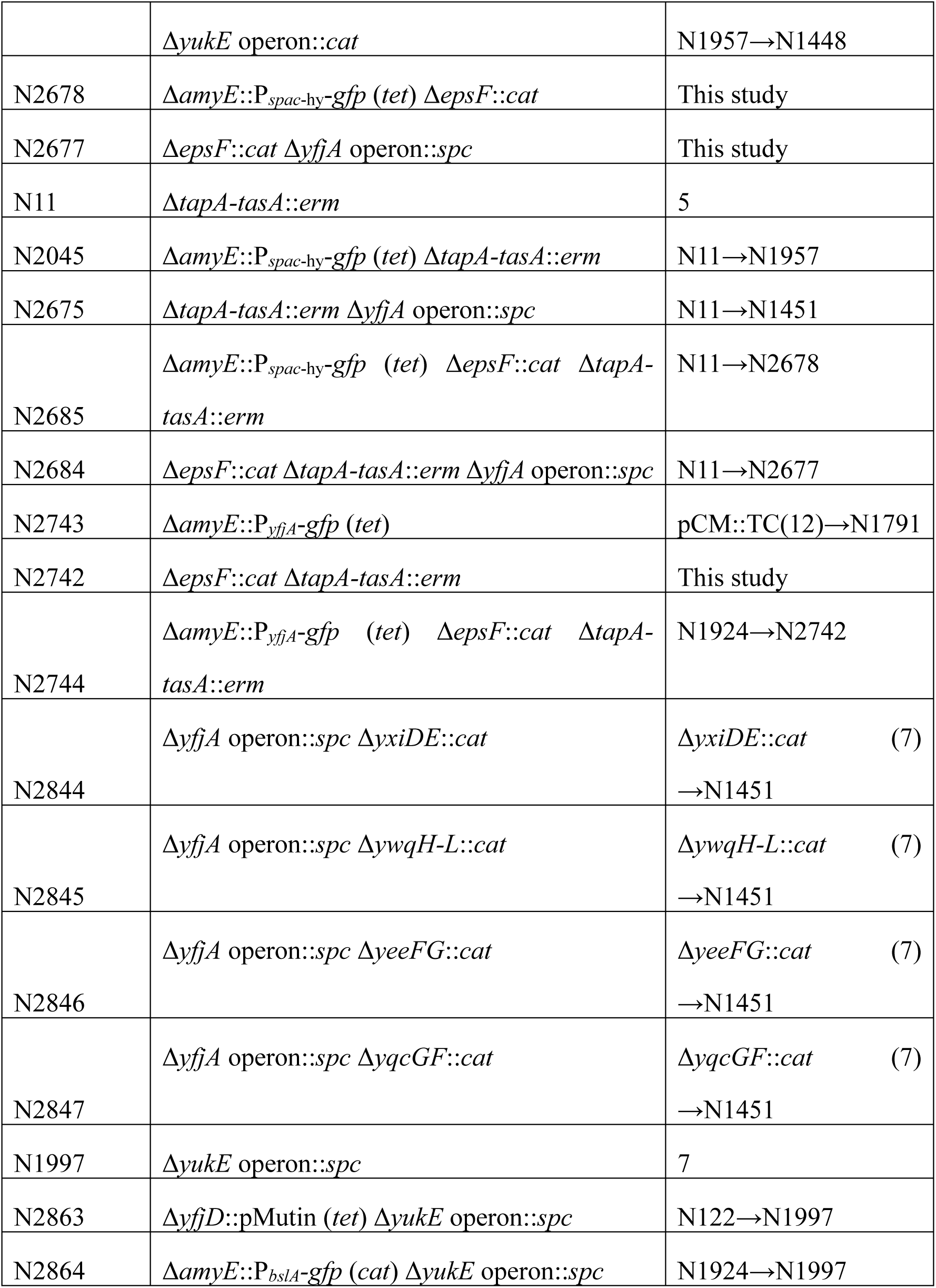
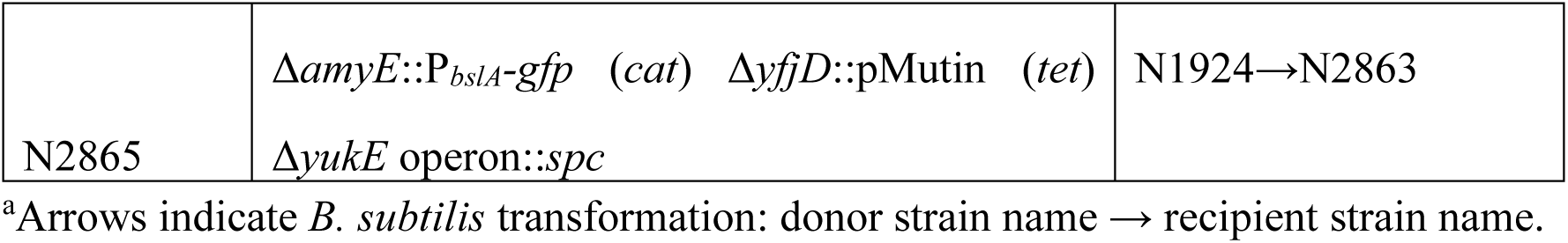
*B. subtilis* NCIB 3610 derivative strains used in this study.

**Table S2.**
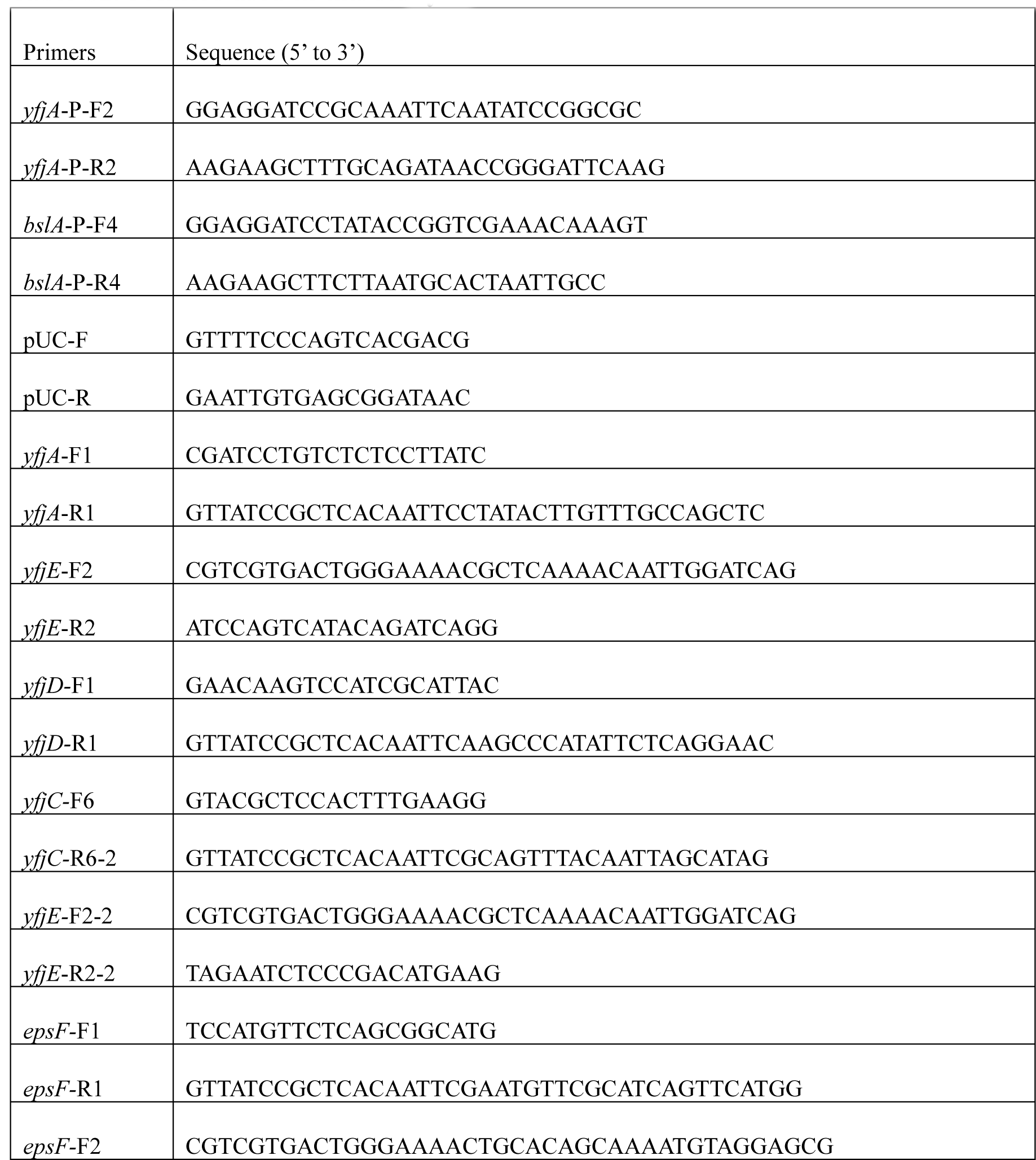
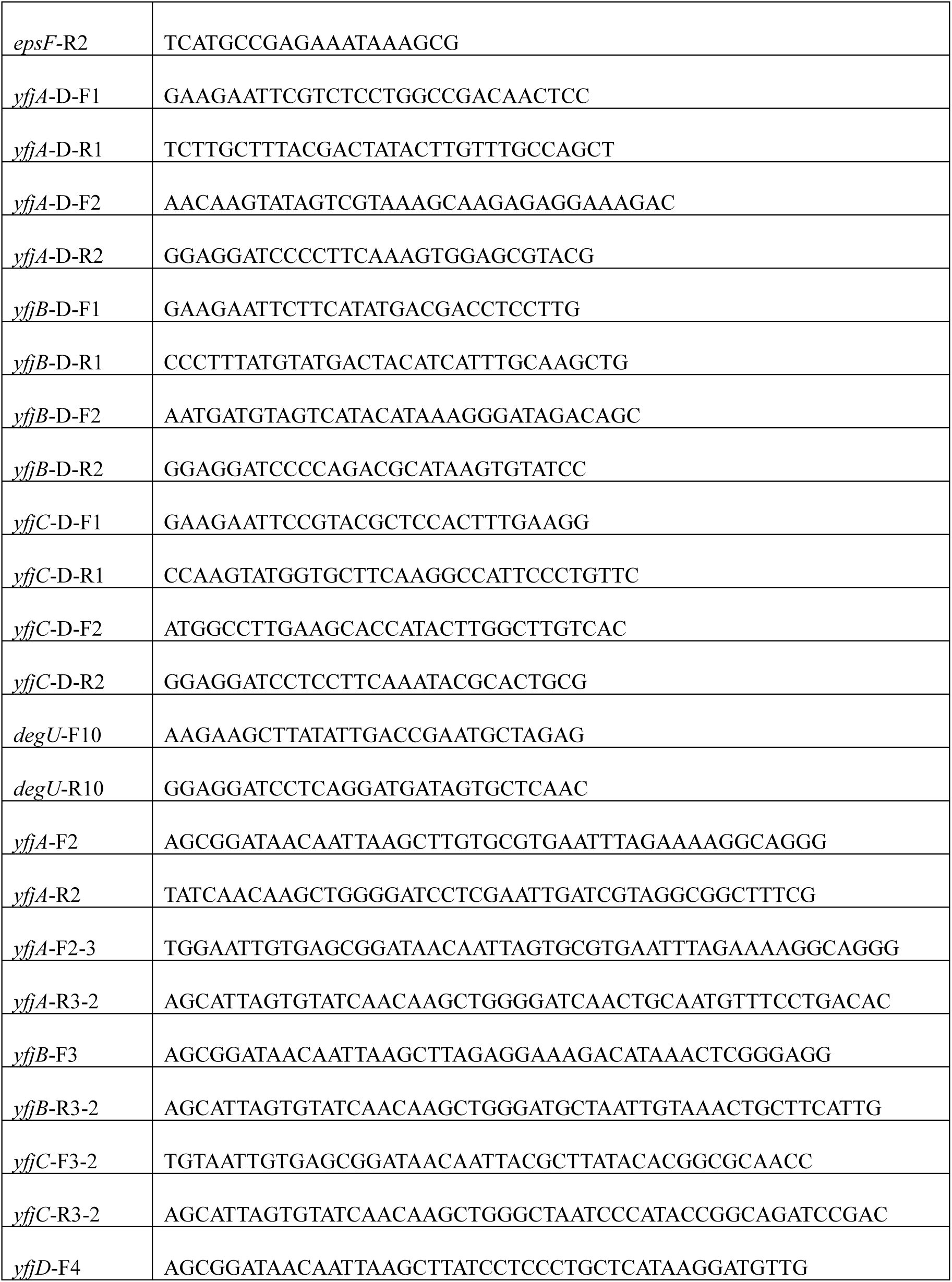
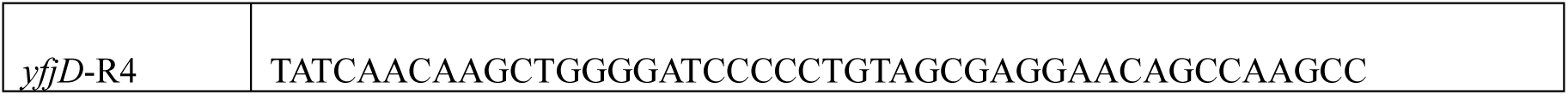
Primers used in this study.

## Appendix figure legends

**Figure S1.**
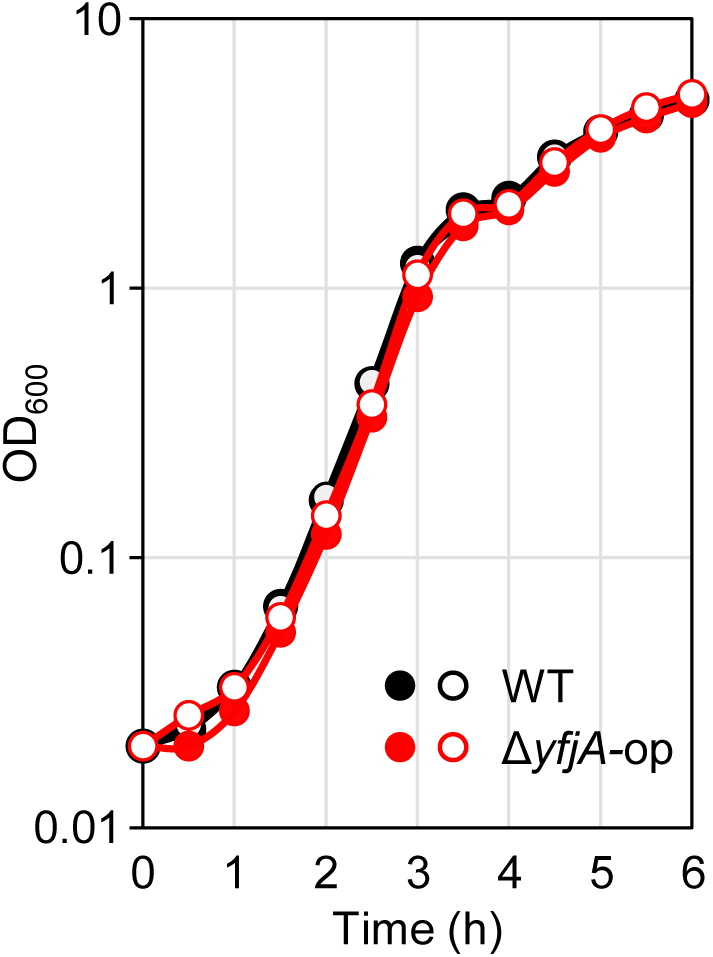
Growth curves of WT and Δ*yfjA-*op strains. Two strains were grown at 37°C in 2×SGG with vigorous shaking. Two independent data for each strain were shown.

**Figure S2.**
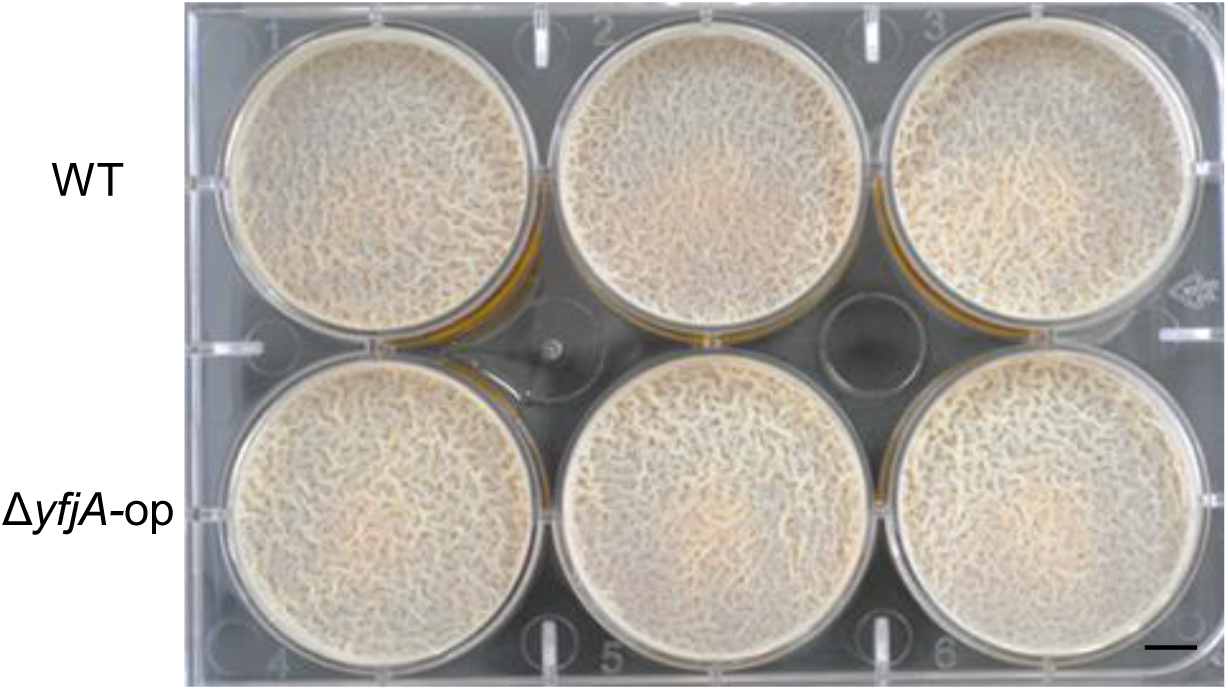
Pellicle formation by WT and Δ*yfjA-*op strains. Exponential phase culture (100 µl of OD_600_ of 0.5 culture) was added to 10 ml 2×SGG in a 6-well plate and then the plate was incubated at 30°C for 48 h. Three independent cultures for each strain were shown. Scale bar, 5 mm.

**Figure S3.**
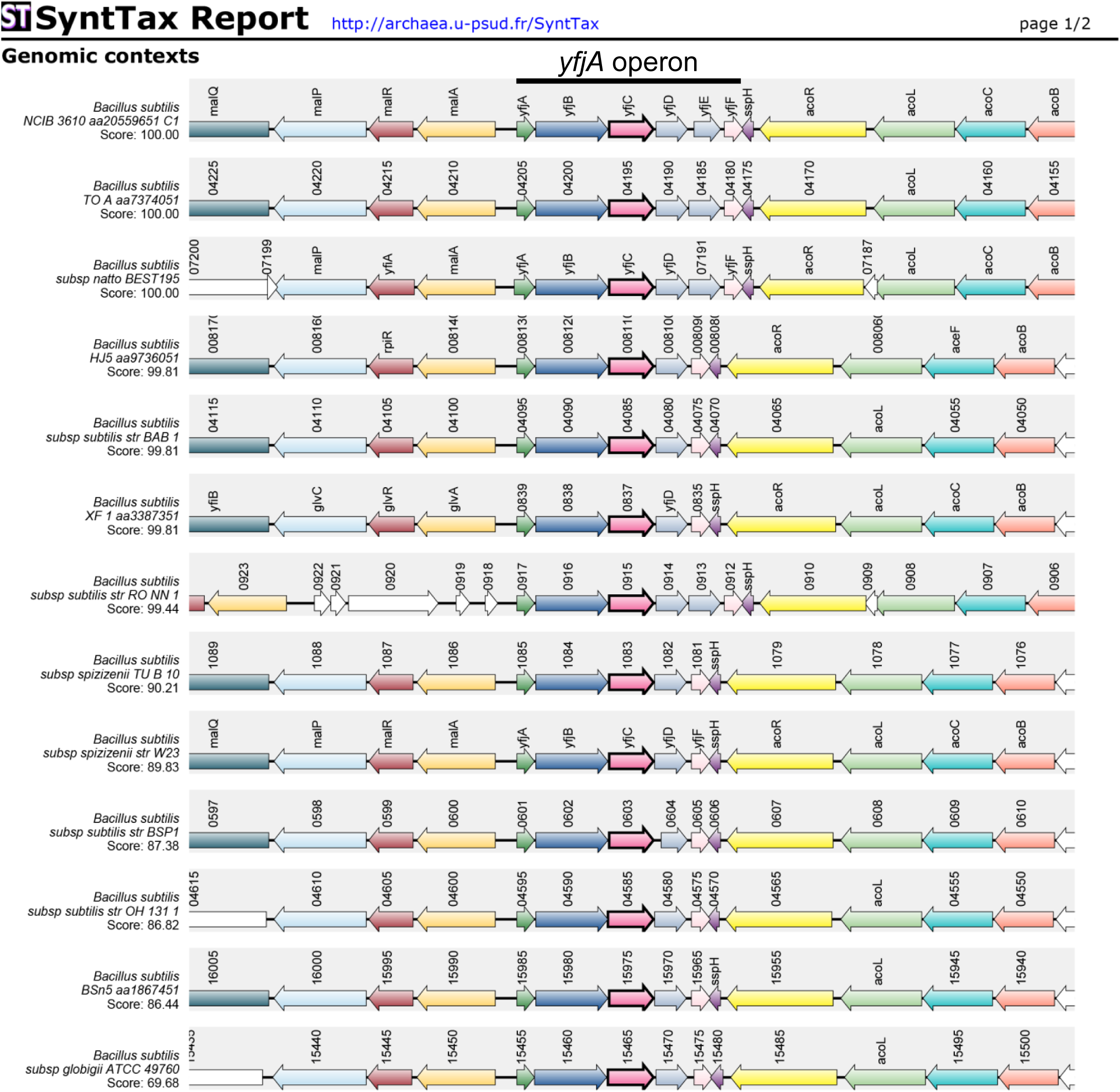
The comparison of the *yfjA* operon region in 13 fully sequenced *B. subtilis* strains. The conservation of the *yfjA* operon in 13 *B. subtilis* strains was analyzed by SyntTax (https://archaea.i2bc.paris-saclay.fr/SyntTax/) using the YfjC sequence as a bait.

**Figure S4.**
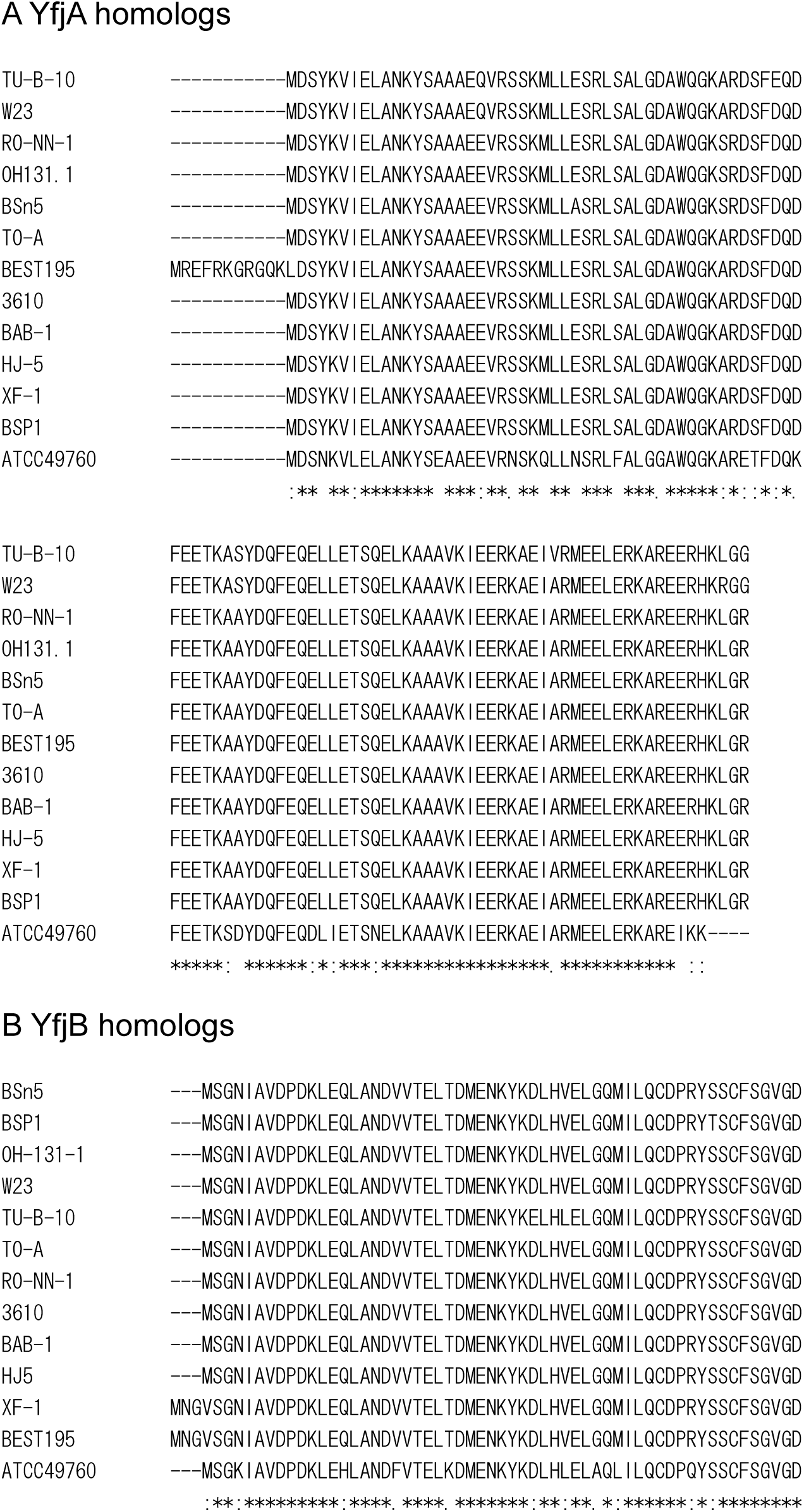
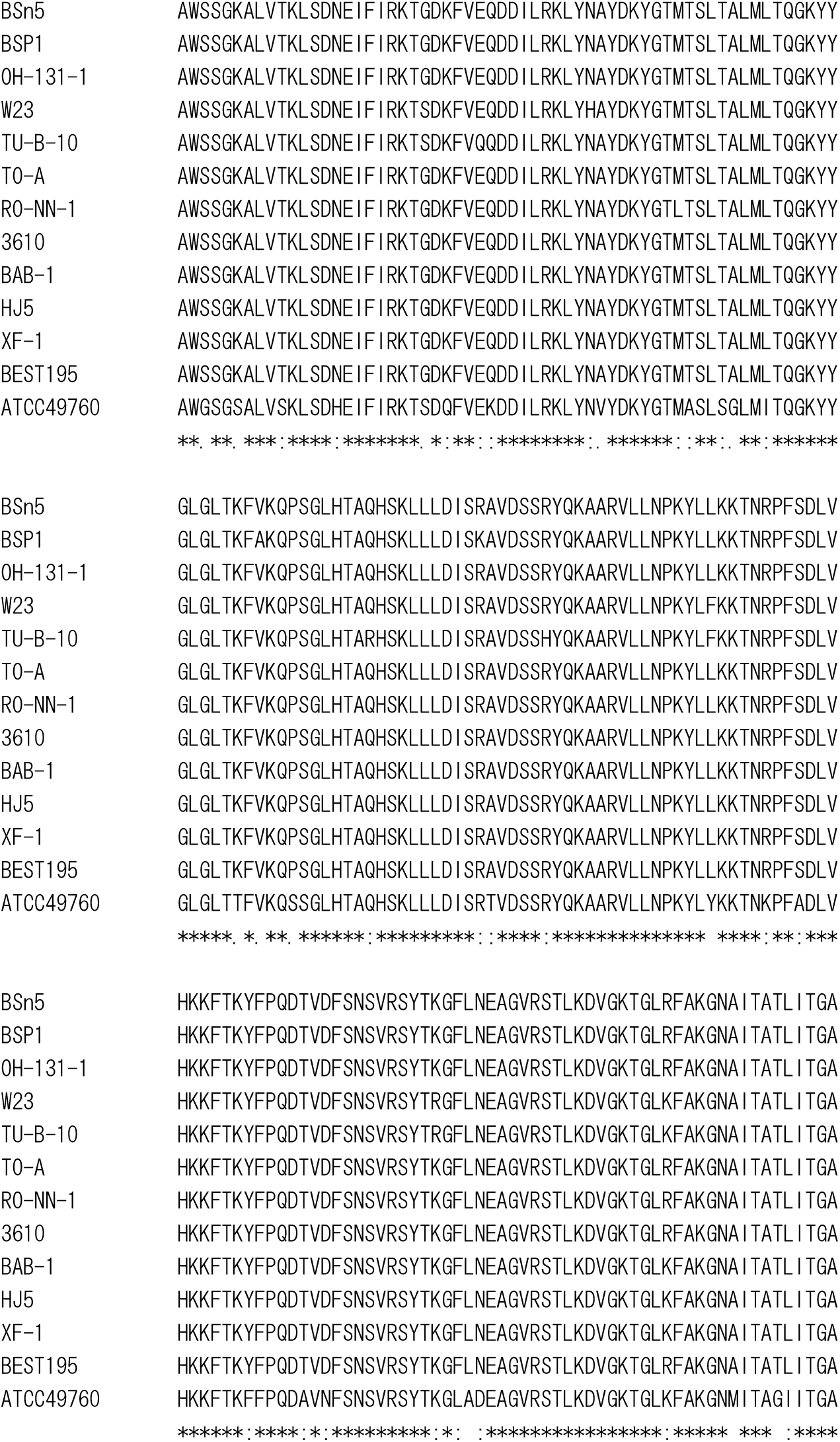
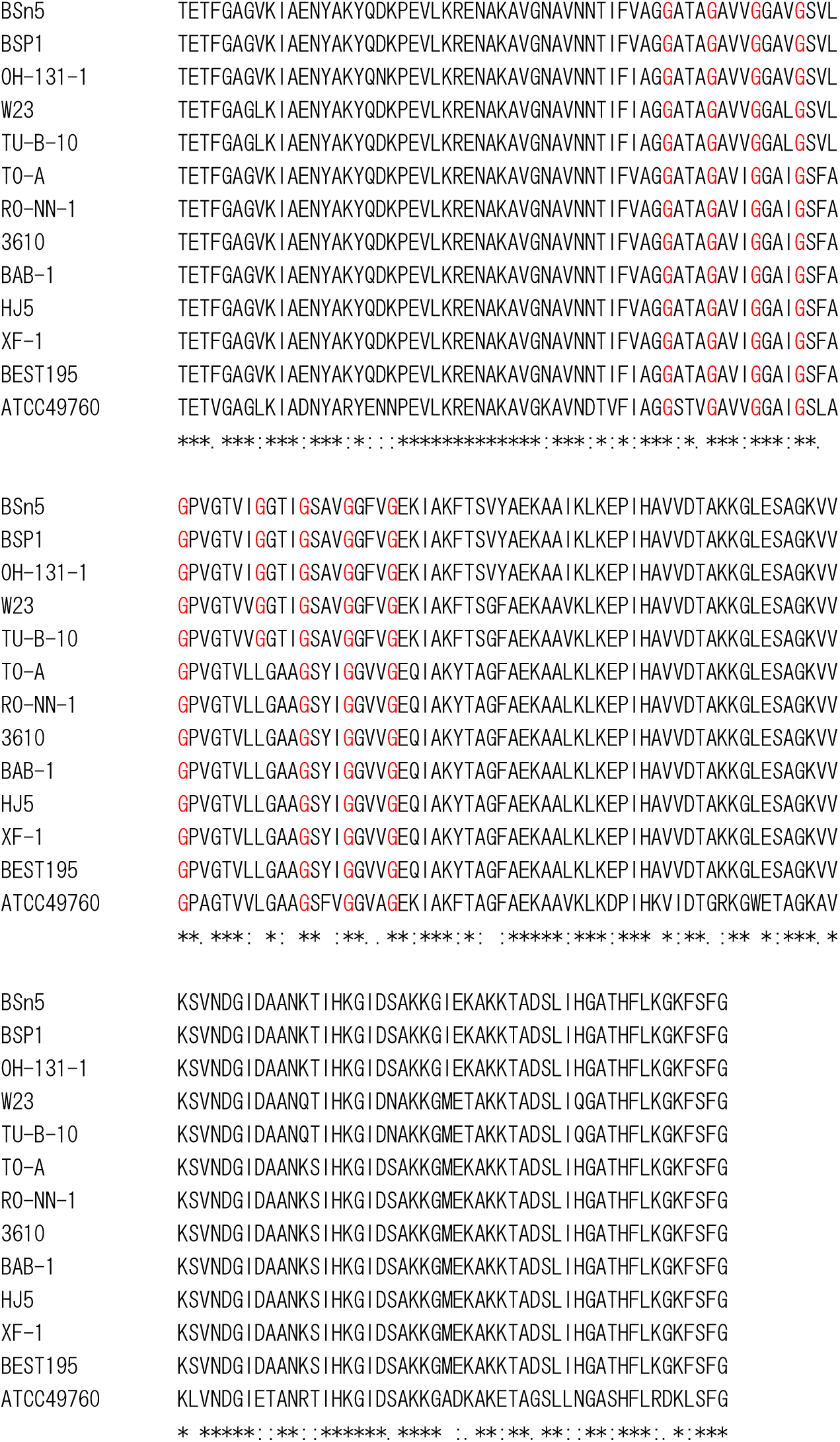
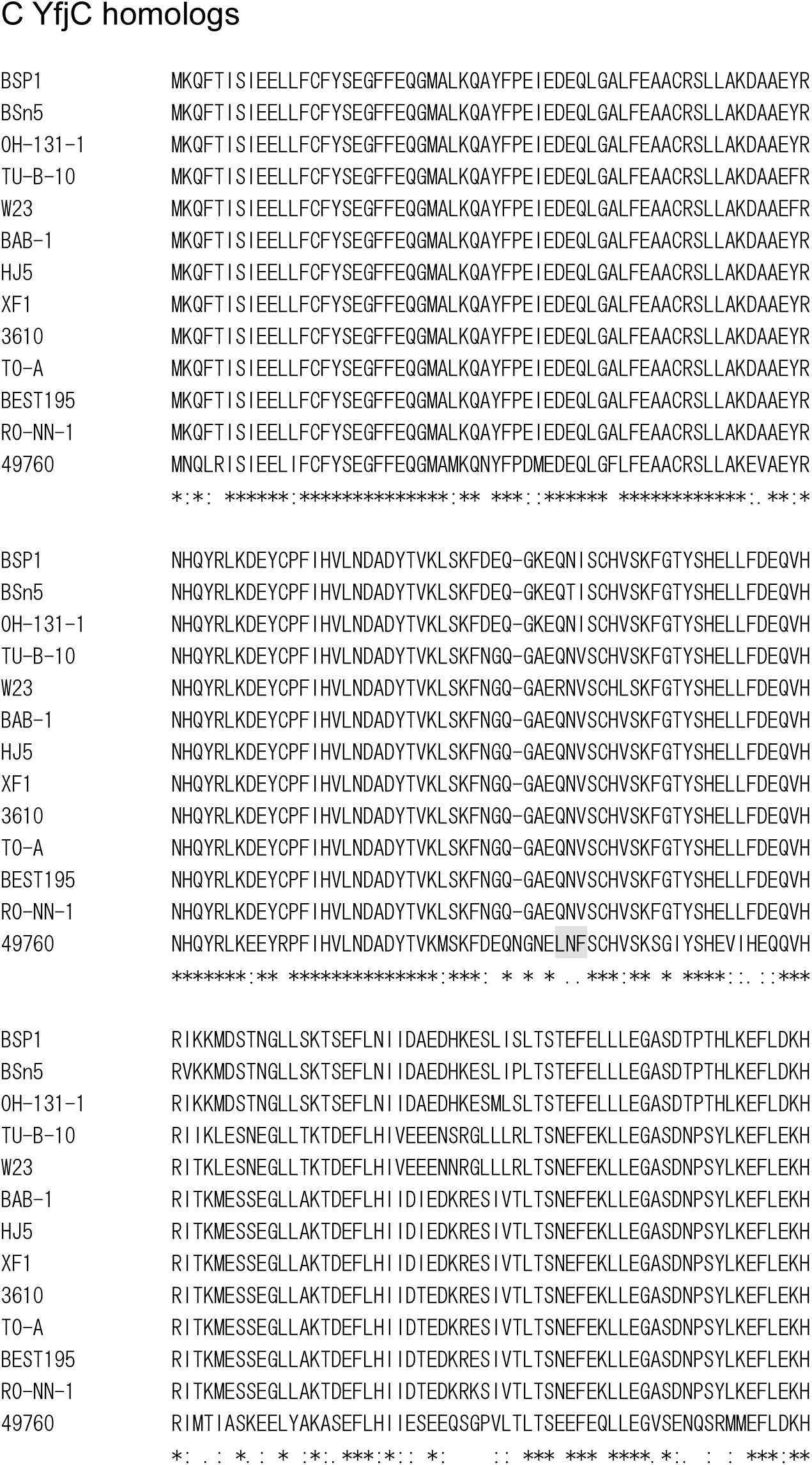
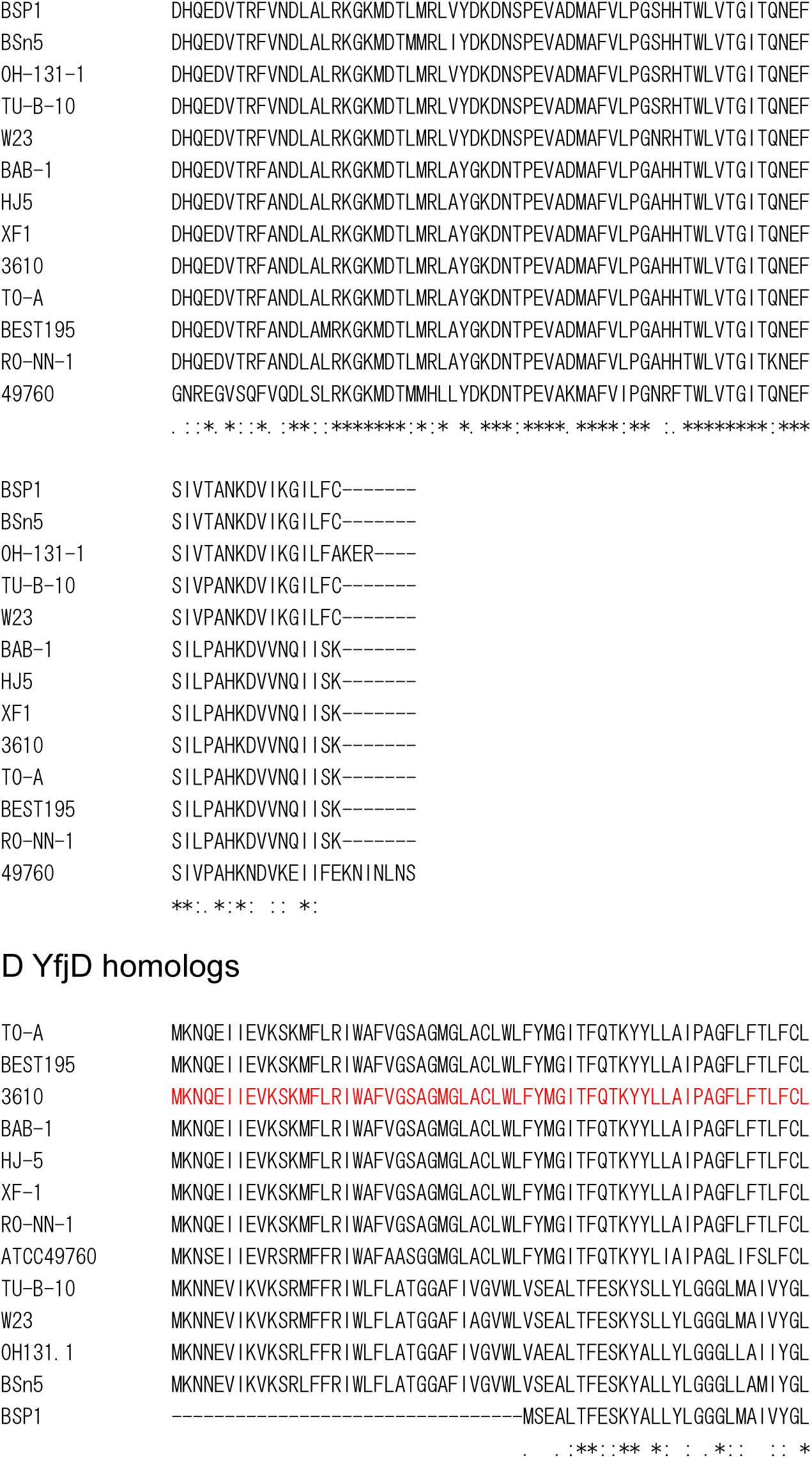
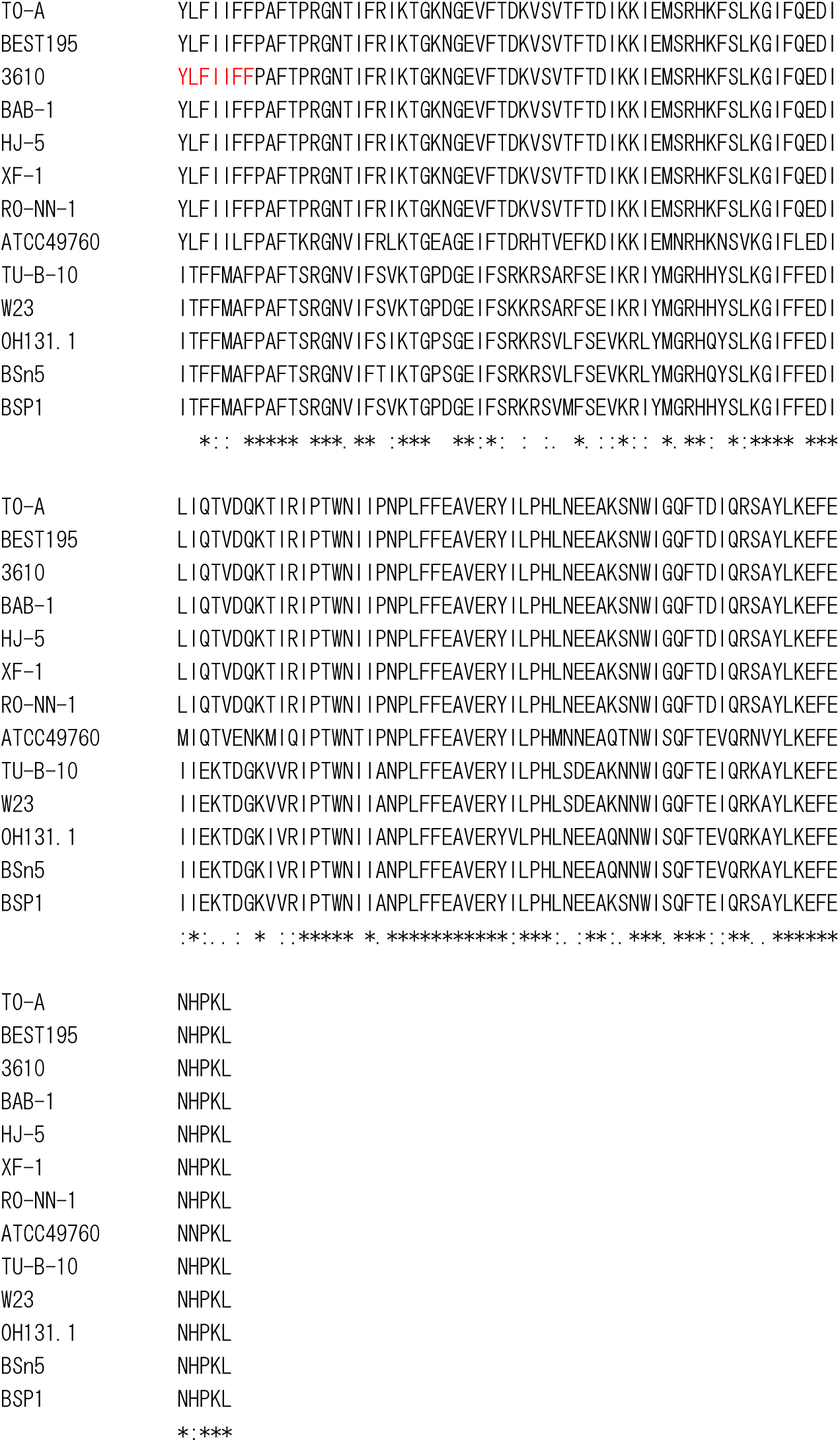
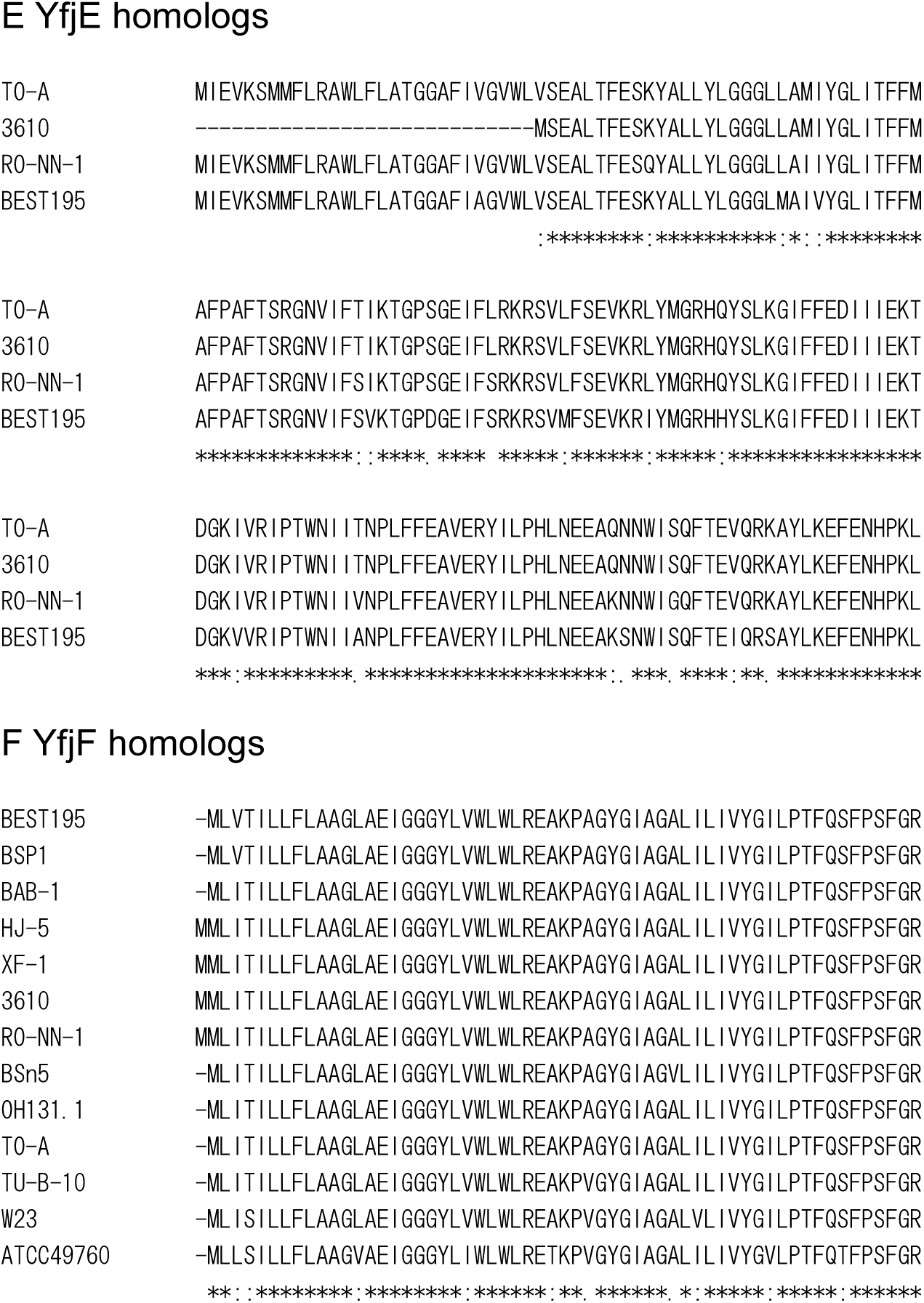
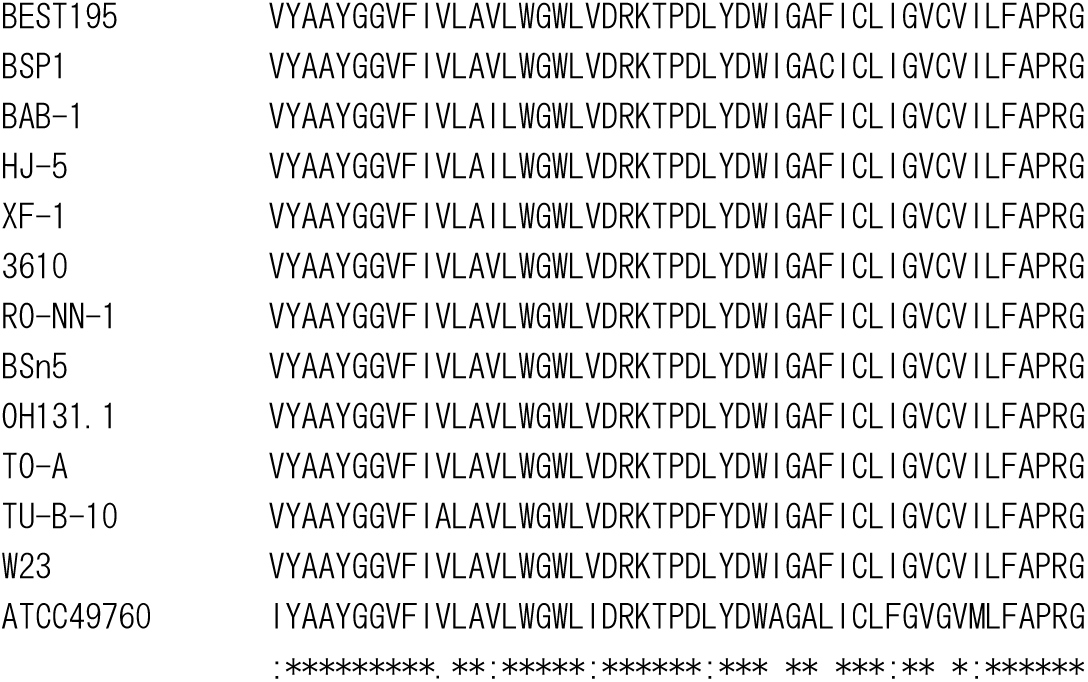
Multiple alignment of *yfjA* operon proteins. A. YfjA. B. YfjB. Gly-xxx-Gly-xxx-Gly sequences are indicated by red G letters in the amino acid sequence. C. YfjC D. YfjD. The signal sequence and transmembrane region of NCIB3610 YfjD is indicated by red. E. YfjE. F. YfjF.

**Figure S5.**
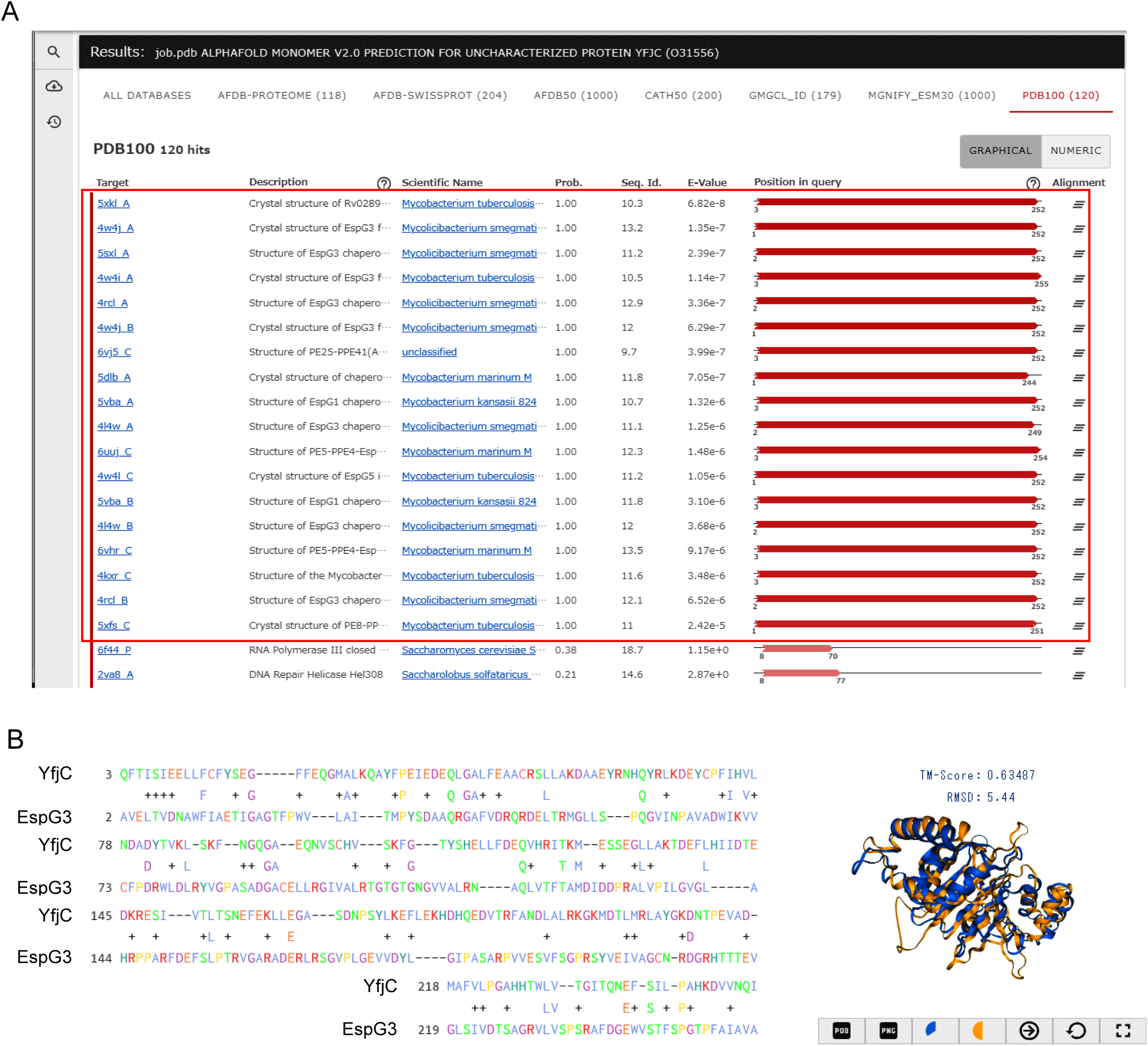
YfjC is structurally similar to EspG chaperons of the mycobacterial T7SS. A. Proteins structurally similar to YfjC were identified by Foldseek search (https://search.foldseek.com/search?accession=O31556&source=AlphaFoldDB). The search result against the PDB database is shown. Identified EspG proteins are boxed in red. B. Comparison of YfjC and EspG3 of *M. tuberculosis* (5XKL in PDB), top listed by Foldseek search in the panel A. Sequence and structure alignments were constructed by Foldseek. Colors in the 3-D alignment: YfjC, blue; EspG3, orange.

**Figure S6.**
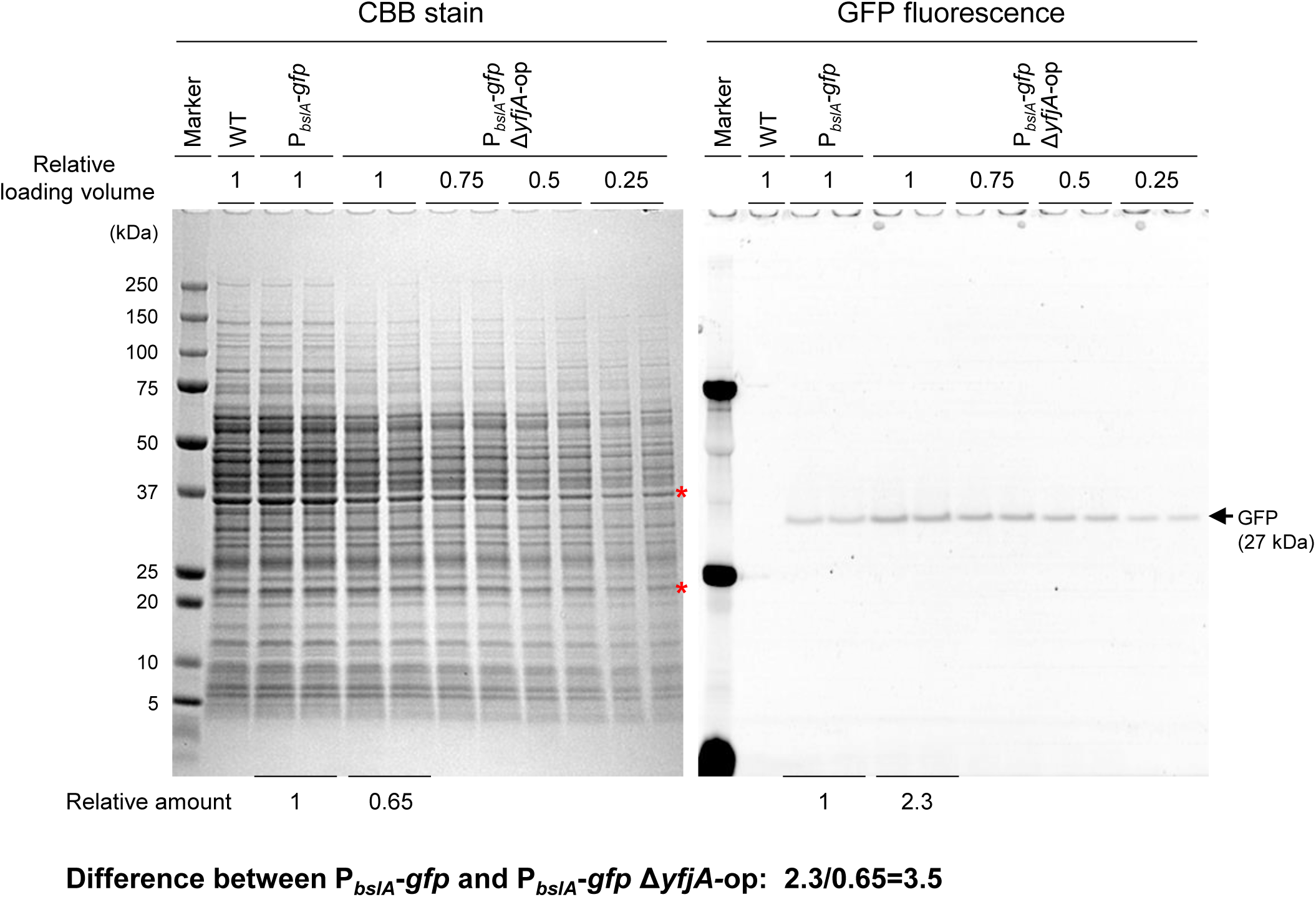
GFP levels in P*_bslA_-gfp* and Δ*yfjA-*op P*_bslA_-gfp* strains. Two-independent crude lysates of P*_bslA_-gfp* and Δ*yfjA-*op P*_bslA_-gfp* strains were prepared from 24 h-old colony biofilms. To quantitatively analyze GFP levels, the undiluted and diluted lysates were loaded on SDS-PAGE, and fluorescence of GFP bands in the gel was detected using a fluorescence imaging system (the right panel). The gel was then stained with CBB (the left panel). GFP and protein were quantitatively analyzed by Evolution Capt (Vilber). Protein bands used for quantitative analysis was indicated by red asterisks. Relative amounts of GFP and proteins were indicated below the gel images.

**Figure S7.**
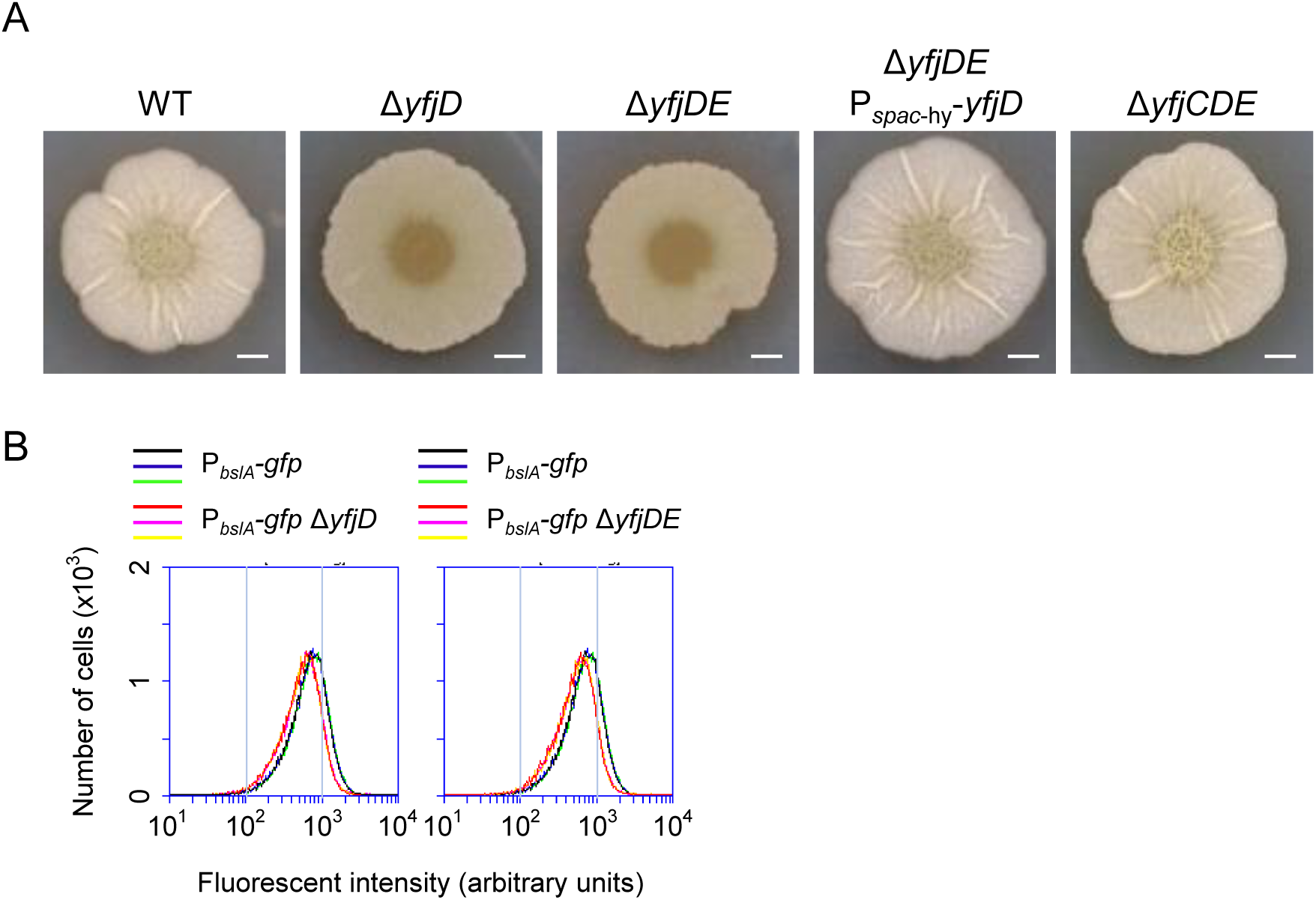
The phenotype of Δ*yfjD* in MSgg medium. A. Colony morphology of Δ*yfjD* and its derivative mutants. Strains were grown for 72 h in MSgg medium. The Δ*yfjDE* P*_spac_*_-hy_-*yfjD* strain was grown in MSgg supplemented with 10 µM IPTG. Scale bar, 2mm. B. Δ*yfjD* and Δ*yfjDE* mutants harboring the P*_bslA_-gfp* reporter were grown in MSgg, and *bslA* expression was measured at 24 h by flow cytometry.

**Figure S8.**
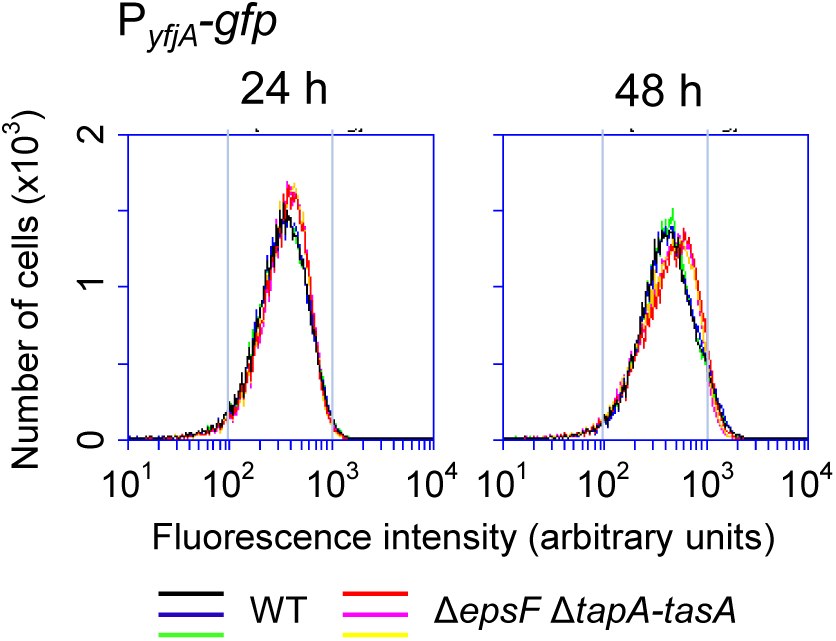
Δ*epsF* and Δ*tapA-tasA* mutations do not prevent expression of the *yfjA* operon. WT and Δ*epsF* Δ*tapA-tasA* strains harboring the P*_yfjA_-gfp* reporter were grown in 2×SGG, and *yfjA* expression was measured by flow cytometry.

